# Invasiveness of Cancer Populations in a Two-dimensional Percolation cluster: a Stochastic Mathematical Approach

**DOI:** 10.1101/2022.03.12.484105

**Authors:** Renlong Yang, Yuanzhi Shao, Chongming Jiang

## Abstract

A framework for the software Unstructured Reaction-Diffusion Master Equation (URDME) was developed. A mitogenic paracrine signaling pathway was introduced phenomenologically to show how cells cooperate with one another. We modeled the emerging Allee effect using low seeding density culture (LSDC) assays to fit the model parameters. Finite time scaling (FTS) was found to be a useful tool for quantifying invasiveness in cancer populations. Through simulation, we analyzed the growth-migration dynamics of BT474 cancer cell populations in-vitro in a 2D percolation cluster and calculated the SPR (successful penetration rate). By analyzing the temporal trajectories of the SPR, we could determine the critical exponents of the critical SPR scaling relation 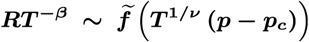. Moreover, the SPR transition point defined according to the FTS theory, ***P***_***c***_, is negatively correlated with the invasiveness of cancer cell populations. The ***P***_***c***_ of the three virtual cancer populations distinctly designated by varying the parameter set of the same model are 0.3408, 0.3675, and 0.4454, respectively. Therefore, the FTS algorithm may be useful in determining invasiveness. Through the simplistic phenomenological paracrine model, inter-cell cooperation and mutual mitogenic boosting are enabled, causing the Allee effect to occur. Such a method could be applied to other circumstances as an example of the quantitatively falsifiable emerging theory.

## 1 Introduction

Invasion of malignant tumors emerges from the interaction between individual cells and the environment, as it is a collective phenomenon. So far, the underlying principles have still not been thoroughly understood.

Paget (1889) has proposed the ‘seed-soil’ hypothesis to explain the organ-specificity of cancer metastasis, marking the beginning of a journey of query spanning more than a century. Jiang et al (2014, 2016) have computed the diffusion exponent of cancer cell population in 2D percolation cluster to evaluate the invasiveness quantitatively. However, the Logistic growth model they used proved inaccurate when cell density is low (Johnson et al, 2019; Korolev et al, 2014; Neufeld et al, 2017). Liang et al (2019) have tried to analyze such invasiveness theoretically and introduced the Allee effect for higher accuracy, and they proved theoretically that the Logistic model and the Allee model behave differently. They also calculated the critical exponents of corresponding growth-migration processes with the finite size scaling (FSS) method, using the successful penetration rate (SPR) as the order parameter. However, their model still needs improvement in assessing parameter uncertainty.

In the studies of (Jiang et al, 2014, 2016) and (Liang et al, 2019), the used parameters were cited from various references and thus lacked consistency. As a result, uncertainty assessment of the parameters could be hindered. Moreover, the spatiotemporal mean-field approximations used in partial differential equations (PDE) in (Liang et al, 2019) could neglect some fluctuation details, especially under the circumstances with low cell densities featuring the Allee effect of concern. Simpson et al (2010) have studied different mesoscopic migration patterns and concluded that formal continuous description might not exist for dynamic systems with strong mesoscopic patterns. Recently, Jin et al (2020) have compared discrete stochastic models and the corresponding continuum limits and pointed out that PDE could not always provide sufficient approximation that discrete models do. However, PDEs under their continuum limits could give a decent depiction of mean behaviors under some certain parameter choices. And they ascribed such deviation to the failure of the mean-field approximations. Therefore, the accuracy of the mean-field approximation is doubtful in research of the growth-migration processes with the emerging Allee effect. To tackle the two difficulties above, it entails a basic mathematical model with higher spatial precision to address the Allee effect and data within a unified experimental framework to assess parameter uncertainty.

The Allee effect emerges from many biological systems, e.g., animal swarms and cell populations. It is a universal and fantastic self-organizing phenomenon (Dai et al, 2012; Frick et al, 2010; Johnson et al, 2006; Myers et al, 2007; Ratzke and Gore, 2016; Redding et al, 2019; Volkov et al, 2007; Wilson et al, 2017). Many similarities have been observed inside these systems, such as identifying a growth threshold, survival through cooperation, or deviation from the Logistic model related to cell densities. These findings suggest a common underlying principle (Angulo et al, 2018; Dennis et al, 2016; Hutchings, 2015; Luque et al, 2016; Maciel and Lutscher, 2015; Sun, 2016). Theoretical development following this lead has been classified into two categories: agent-based models that use few, simplistic interaction rules to depict migration and the life cycles directly (Boettger et al, 2015; Colon et al, 2015; Fadai et al, 2020; Castner et al, 2011; Gilroy and Lockwood, 2016; Goto et al, 2020; Manor and Shnerb, 2008; Mendonc;a and Gevorgyan, 2017; Murphy and Johnson, 2015; Pires and Duarte Queiros, 2019; Ribeiro, 2015), and a more macroscopic approach in the cell density limit, relying on sophisticated PDE terms designated for the system under concern (Gao et al, 2016; Korolev, 2015; Mendez et al, 2019; Meyer et al, 2018; Neufeld et al, 2017; Newman et al, 2004). Huge parameter spaces and a myriad of behavioral patterns are common characteristics among all these models. The main obstacles to understanding the underlying principles are the complexity of the interactions in existing experimental settings and limited parameter control. Johnson et al (2019) have performed relatively accurate parameter fitting of the weak Allee stochastic models to the low seeding density culture LSDC assays. However, only the principles of the temporal evolution of the total cell number were summarized with no further surveys on the evolution of the spatial distribution.

The existence of an unknown mitogenic paracrine signaling pathway was assumed in this study to represent intercell cooperation. It was used to interpret the phenomenological mechanism underlying the emergence of the Allee effect. The spatial accuracy of the description was enhanced through a discrete Poisson process model. The mitogenic role of paracrine signaling pathways can be seen in (Alawin et al, 2017; Castillo et al, 2017; Khalilpourfarshbafi et al, 2017; Li, 2019; Xu et al, 2019). This study finds a negative correlation between the 2D percolation cluster invasiveness and the percolation transition point *p*_*c*_. The introduced paracrine-Allee effect spatiotemporal Growth-Migration system is not only applicable to the analysis of cancer population invasion but also sheds new light on the emerging phenomena related to a large category of subjects, including social animal swarm dynamics (Frick et al, 2010; Myers et al, 2007; Redding et al, 2019; Volkov et al, 2007) and the collective dynamics of microbial cultures (Keeling et al, 2003; Ratzke and Gore, 2016).

## 2 Model Construction

The introduction has called for a more accurate depiction of the spatiotemporal dynamics of cancer cell populations in a 2D percolation cluster. A spatial description of the Allee effect was necessarily introduced to address this. Besides, it should also be considered whether the parameters can be fitted within a unified experimental framework.

To achieve these two ends, the following requirements should be satisfied:

1. Addressing the present imbalance between theory and experiment in cell population biology and helping devising experiment settings to enable uncertainty assessment of Max Likelihood Estimation (MLE).
2. Getting rid of the constraints of the mean-field hypotheses in continuum models and changing for stochastic models and using the moment closure method (Fisher, 1922; Froehlich et al, 2016; Johnson et al, 2019; Raue et al, 2009, 2013) to construct the likelihood in the MLE procedure.
3. Flexible, customized coarse-graining in the regions of less interest for saving computational resources. In other words, achieving high spatial precision (∼ 50 *μm)* computation using dense mesh where needed and keeping flexible implementation of less dense mesh in the rest of the regions.
4. Scalability. Convenient extension to the stochastic description of chromosome mutation/modification in individual cells, enabling continuous, sustainable model evolution.

Representative cancer models over the past two decades were surveyed. Following Table 1 is a summary on how they meet the requirements above.

**Table 1.**
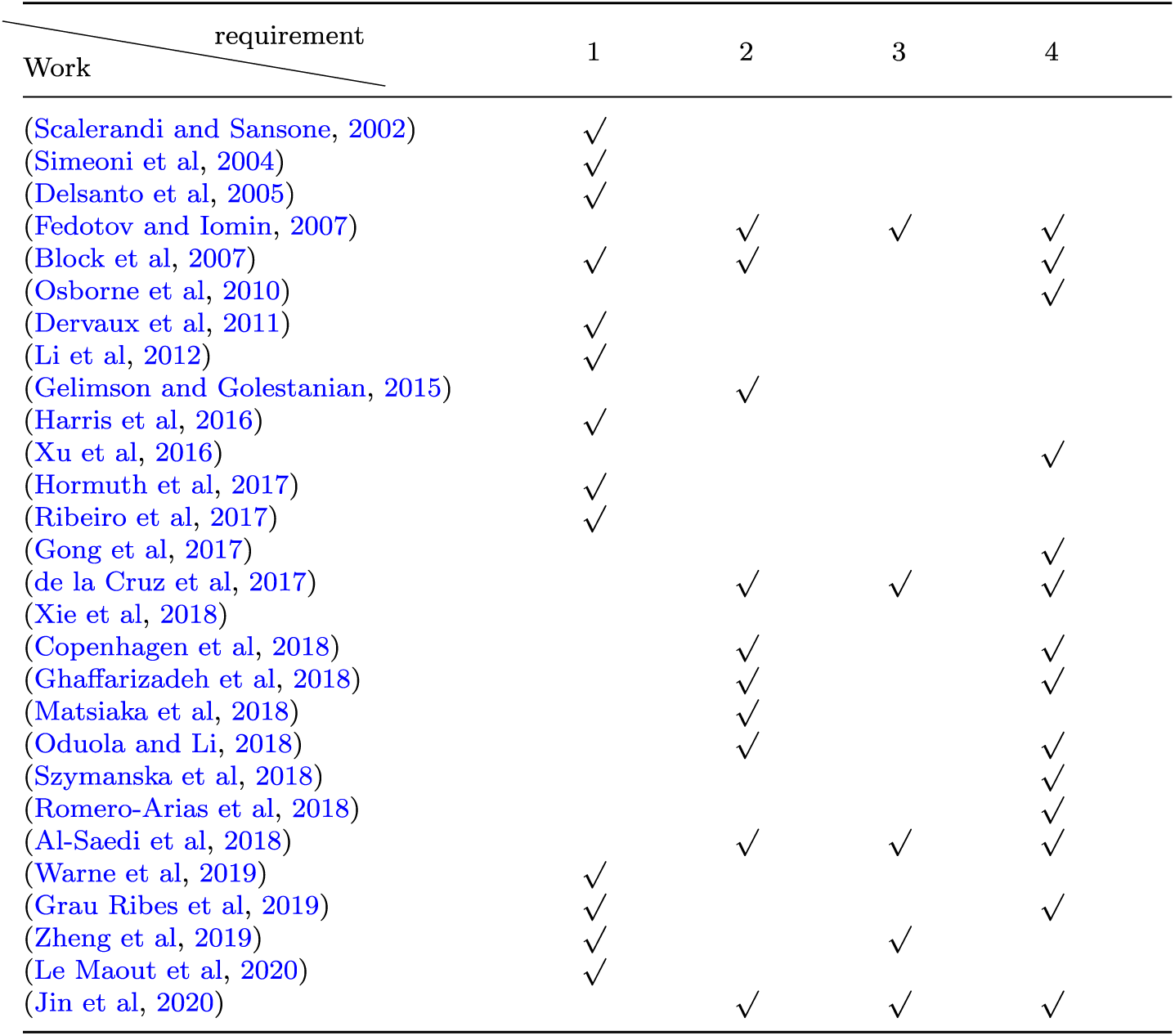
Summary of cancer modelling in the past two decades

The studies above simply excel in one way or another, but they cannot meet all the requirements simultaneously. Engblom (2019) have proposed to use the Unstructured Reaction-Diffusion Master Equation (URDME) method (Drawert et al, 2012) to address the two spatiotemporal scales of both individual cells and population in the very same paradigm, thus explaining qualitatively how periodic spots at the population scale emerged from the Notch signaling pathway at the cellular scale. Such a paradigm meets all the four requirements above. In light of this, URDME was used within the experimental framework (Johnson et al, 2019) for modeling and MLE assessment. Finally, whether the finite time scaling (FTS) method could be useful for quantitative invasiveness measurement was analyzed. The reason for the choice of the experimental framework is that Johnson et al (2019) have reported and explained the emerging Allee effect in LSDC assays.

The experiments in (Johnson et al, 2019) were carried out in planar space, the cancer cells were seeded initially, and the media was changed regularly. It was observed at a low cell density that the weak Allee effect could be revealed by fitting the stochastic models to LSDC data. The stochastic computational method Johnson et al (2019) have used enables backtracking of the underlying phenomenological local interactions through the emergence of the Allee effect. Such interactions include the proliferation and death of individual cells. The method could also enable the moment closure method (Fisher, 1922; Froehlich et al, 2016; Johnson et al, 2019; Raue et al, 2009, 2013) for MLE. The moment closure method has been made plausible with the help of the model’s stochasticity and enormous experimental ensemble data obtained.

It is urged that the Allee effect should be depicted at a higher spatial precision (∼50 *µm*) because the Allee effect calls for addressing low cell density circumstances. This was achieved by customizing the triangular mesh of the domain. More details on the mesh design can be seen in Section 5.2 and Section S6 in the Supplementary Material. The paracrine mitogenic signaling pathways were introduced as the Allee effect mechanism. More specifically, it is assumed that secretion, diffusion, the capture of the inter-cell signaling molecules should be related to the mechanism underlying the emergence of the Allee effect. Subsequently, model parameters corresponding to those assumptions were designated and fitted.

After the following report on how the spatial depiction integrating the Allee effect was designed and fitted, the focus will then be brought back to the question brought about by the seed-soil theory mentioned in the Introduction: how to measure the invasiveness of cancer cell populations in 2D percolation cluster quantitativelypercolation cluster. Table 2 illustrates model settings, definitions, and symbols. For more details, see Section 5.1. The model consists of a series of independent Poisson events, each of which is depicted by a Poisson rate parameter. Following are a series of model assumptions and the corresponding equations for the Poisson events. (Assumption 1)The Unboosted Cells and Boosted Cells are assumed to divide into two daughter cells with the Poisson rate parameters, *η*_2_ and *η*_3_, respectively.

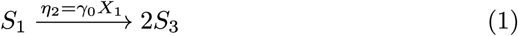

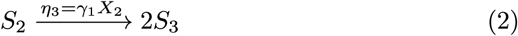

**Table 2:**
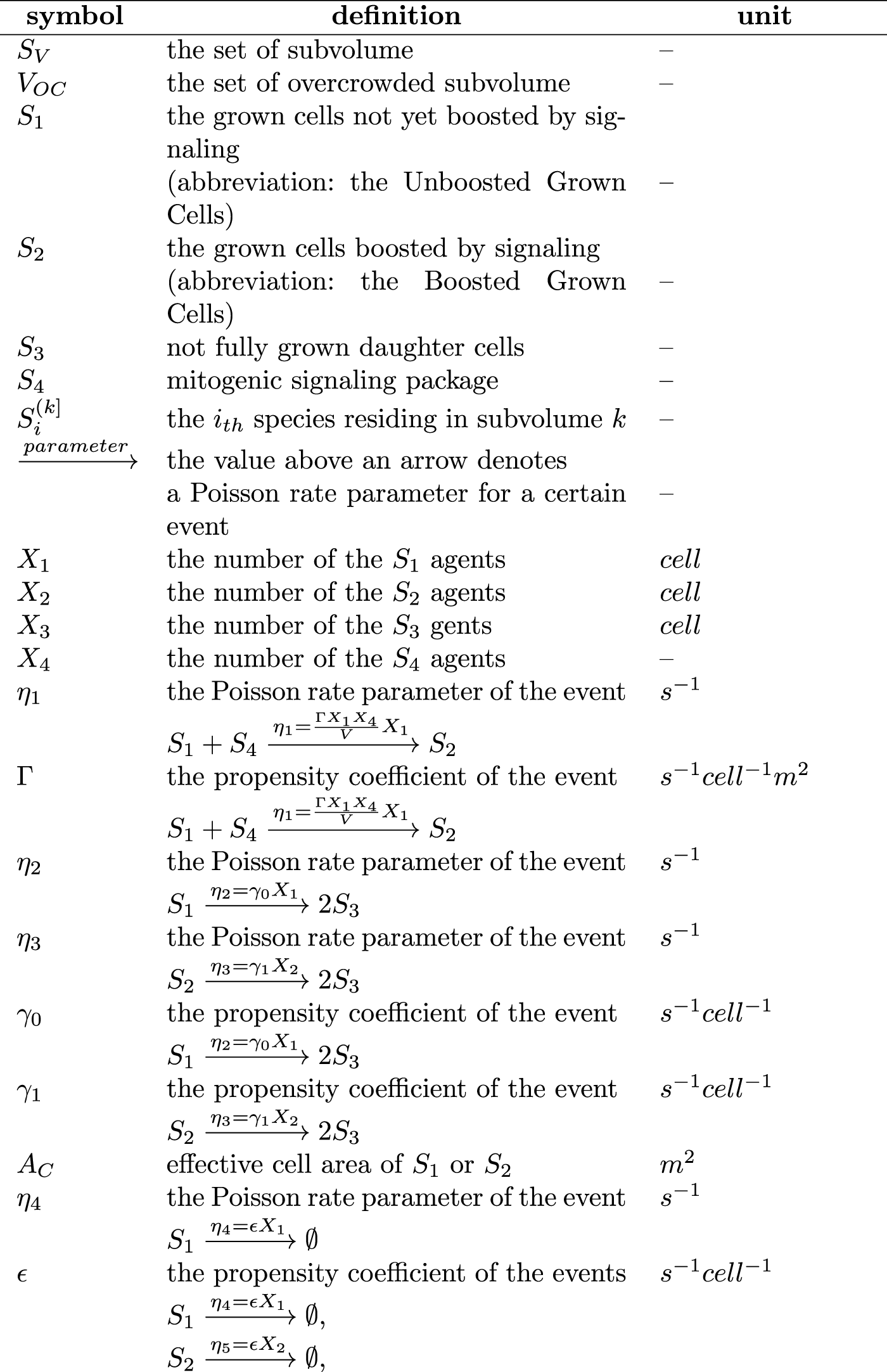
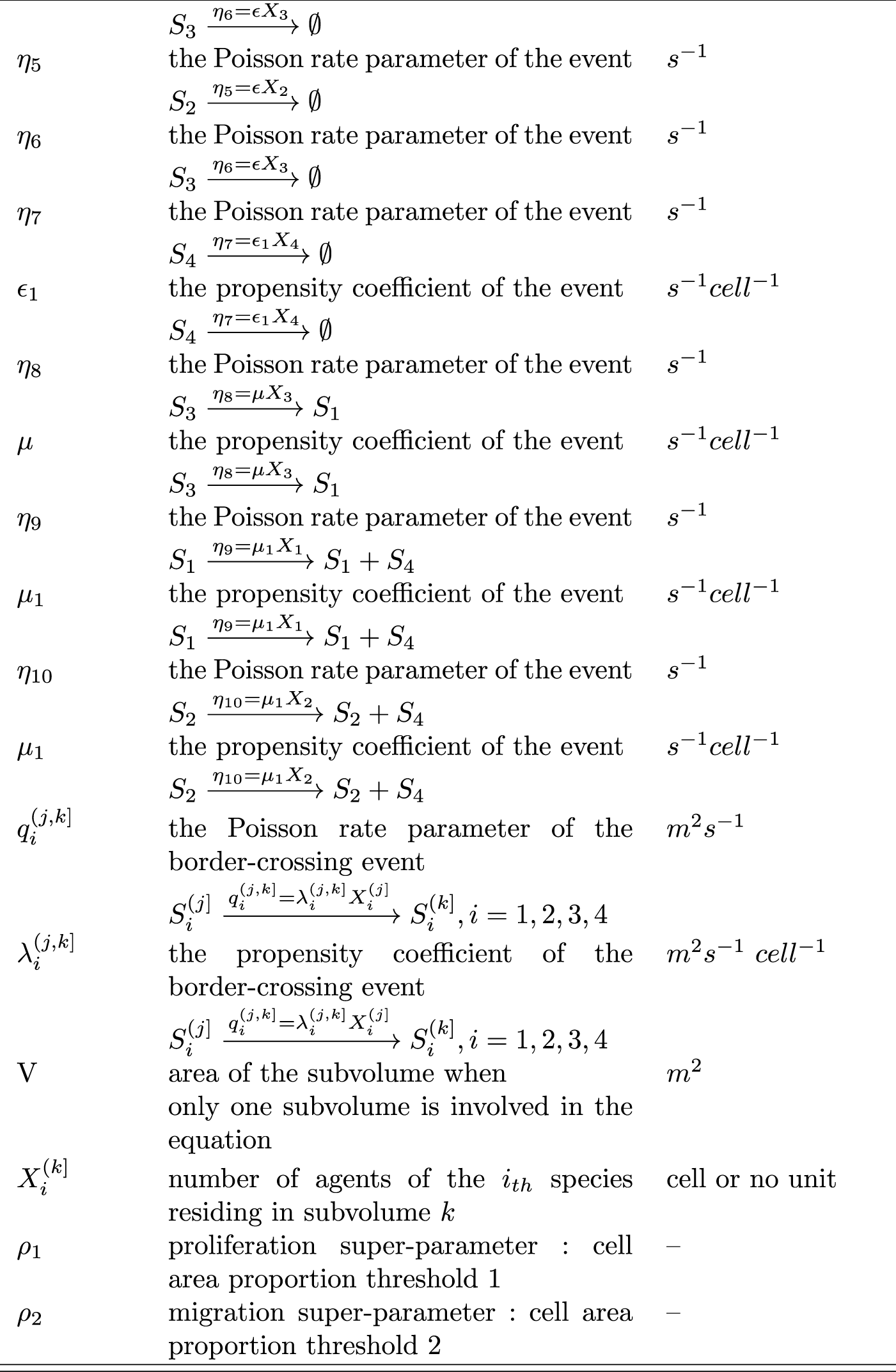
Definition of the symbols

(Assumption 2)It is assumed that the area of the daughter cells is only a half of that of their mother just after division and that the daughter cells develop into fully grown Unboosted Cells with the Poisson rate parameter *η*_8_.

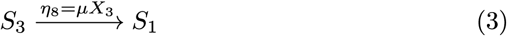

(Assumption 3)A pseudo-particle approximation is introduced to mitigate the computational complexity. A pseudo particle named ‘signaling package’ was introduced, and the interactions between those packages and the cells are called ‘cell-package interactions. The cell-package interactions were designed to take the place of thecell-molecule interactions. Cell-package reaction-diffusion processes are used to describe the secretion, diffusion, and capture of the mitogenic signaling molecules at a greater spatiotemporal scale than that of cell-molecule reaction-diffusion processes. It is assumed that the Unboosted Cells and Boosted Cells release a mitogenic signaling package with the Poisson rate parameter *η*_9_ and *η*_10_, respectively.

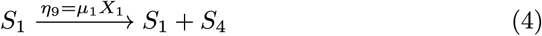

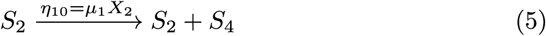

(Assumption 4)It is assumed that the Unboosted Cells capture the mitogenic signaling packages and turn into the Boosted Cells with the Poisson rate parameter *η*_1_.

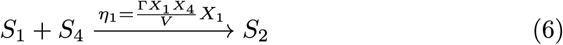

(Assumption 5)It is assumed that the Unboosted cells, the Boosted Cells, and the daughter cells die with the Poisson rate parameters*η*_4_,*η*_5_,and *η*_6_, respectively.

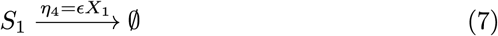

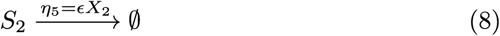

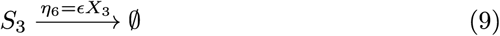

(Assumption 6)It is assumed that the mitogenic signaling packages are decomposed with the Poisson rate parameter *η*_7_.

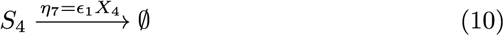

(Assumption 7)The migration of the cells and the transport of the signaling packages are depicted by the border-crossing events. It is assumed that the event ‘an agent of the *i*_*th*_ species moving across the border of the *j*_*th*_ subvolume and into the *k*_*th*_ subvolume’ with the Poisson rate parameters 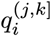.

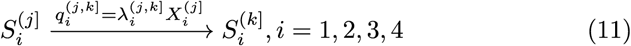

## 3 Analysis and Results

The ‘Achilles’ Heel’ of biological sy stems research is the incapability to control the parameters and the complexity of the interactions. Scale invariance (Lesne and Lagues, 2011) of the subjects could be used to address these difficulties. It is assumed that researchers could take the occupation probability as the control parameter in the future and observe the critical behaviors. (Diao et al, 2019; Kim et al, 2018; Ouyang et al, 2020; Zheng et al, 2019) could already manufacture chambers at the size of 200 1000 µm in artificial ECM, using the 3D-printing technology. Therefore, it is promising that the size of the pores can be contained to several dozen micrometers.

The boundary conditions of the percolation cluster are highly complicated. Unfortunately, the finite size scaling (FSS) analysis method needs a huge number of grid points. Therefore, obtaining the data needed for finite size scaling seems infeasible, judging from the sheer computational complexity. Despite this, SPR is also related to the time scale of the experiment, besides the spatial scale. Furthermore, samples at multiple time scales could be obtained through a single run of an assay/a simulation. Thus, the FTS method, closely related to the time scale, appears to be more interesting. It really makes one wonder whether the critical values derived by the FTS method could be used to depict the invasiveness of the cancer cell populations in a 2D-percolation cluster.

Based on the logic underlying the FSS method, computation of the Growth-Migration process was analyzed with the FTS method proposed by (Sorge, 2018) (see section 5.6). Critical exponents were derived and were designed to also be measured experimentally.

Evolution from Oh to 800h was computed. In the computation, two Unboosted Cells (S1) were seeded in the initial seeding site exactly in the middle of the leftmost column of the 2D percolation cluster. The time trajectories fluctuate remarkably because of the stochasticity and the low seeding number. A considerable number of repeated runs were carried out for results to work with. For each given value of occupation probability *p*, a batch of site configurations was generated. Hence, each site configuration generates a batch of stochastic temporal trajectories. From these statistics, the SPR can be calculated. Further details are provided in the following sections.

### 3.1 The SPR and the uncertainty of computation

A 17×17 square percolation lattice was designed, The edge length of each site is 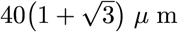 (see section 5.2). See Fig. S1 in Supplementary Material S2 for an illustration of the percolation assay settings. Two Unboosted Cells (*S*_1_) were seeded in the initial seeding site in the middle of the leftmost column. Then they are used in the runs of the temporal trajectories. The event that any cell gets to the rightmost column is called ‘Successful Penetration’ (SP). The offset of ‘Successful Penetration’ was observed and recorded every time step. Given the *j*_*th*_ occupation probability *p*_*j*_, the computation was carried out in the *k*_*th*_ site configuration, and the offset of ‘Successful Penetration’ at the time point *t*_*l*_ was recorded. The “penetration state” is defined as a Boolean value *B*_*jk*_ (*t*_*l*_). Initially, *B*_*jk*_ (0) = 0 and remain zero, and during computation if ‘Successful Penetration’ takes place at *t*_*l*_, then *B*_*jk*_ (*t*_*l*_) = l, ∀*t* ≥ *t*_*l*_.

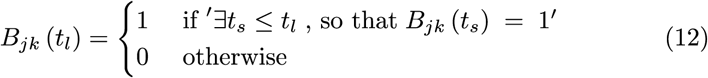

Next, *B*_*j,k*_ (*t*_*l*_) temporal trajectories of 1000 site configurations were averaged to obtain one temporal trajectory of the SPR. The temporal trajectory of the SPR is denoted as *F*(*t*|*p*_*j*_).

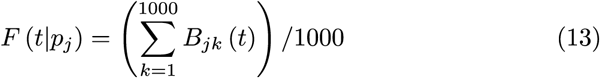

It is shown in Fig. 1 that the uncertainty of F (t IP1) are drawn as bands in the vicinity of the ensemble means *< F* > *F* (*t*|*p*_*j*_). *< F* > *F* (*t*|*p*_*j*_) is obtained by averaging 100 replicas. The ensemble mean

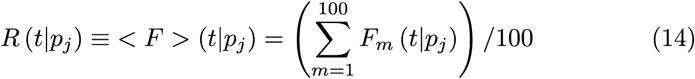

is denoted as *R* (*t*|*p*_*j*_), where *m* represents the *m*_*th*_ replica for the *k*_*th*_ site configuration.

**Fig. 1.**
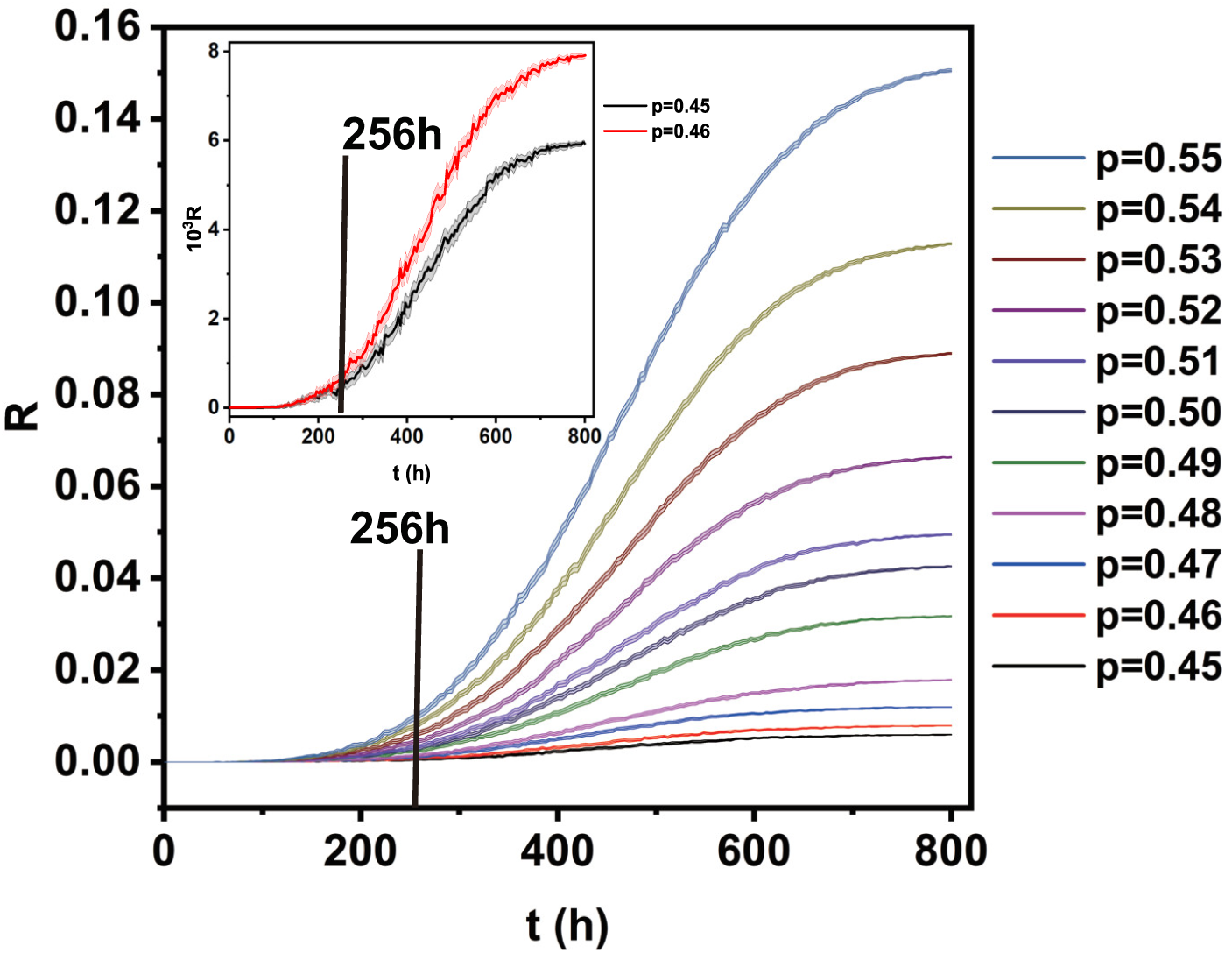
Dependence of the ensemble mean, SPR (denoted as *R)* and time *t* for different occupation probability p.The uncertainty of *F (t*|*p*_*j*_) are drawn as bands in the vicinity of each *R (T*|*p)∼ p* curve.

For *t* >= 256*h*, it can be seen that there is almost no overlapping between the relatively localized uncertainty bands. The low relative uncertainties (all less than 0.2 for *t* ≥ 256*h*) follow the computation of 100 replicas for each configuration. This has guaranteed the reliability of the subsequent FTS analysis.

In the top left corner of Fig. 1 is the magnification of the uncertainty bands for both p = 0.45 and p = 0.46. The two bands can be distinguished from each other at *t* ≥ 256*h*. The ensemble with *p* = 0.45 have the widest uncertainty band and note that all those bands in Fig. 1 were contained in a relatively narrow range.

### 3.2 Determination of the Max Permitted Scale (MPS) *T*_*max*_

After obtaining *R* (*t*|*p*_*j*_), *j* = 1, 2, …, the FTS analysis was carried out for these temporal trajectories. Several finite system time scale *T*_*i*_, and several occupation probabilities *p*_*j*_ within the critical region were set. The discrete data points *R*_*i*_,_*j*_ for a given scale *T*_*i*_ is defined as

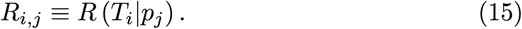

and *T*_*min*_ = 256*h*. The set of occupation probabilities *p* defined in equation (16) will be used in all the FTS procedures throughout this work.

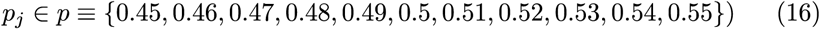

and the time scales

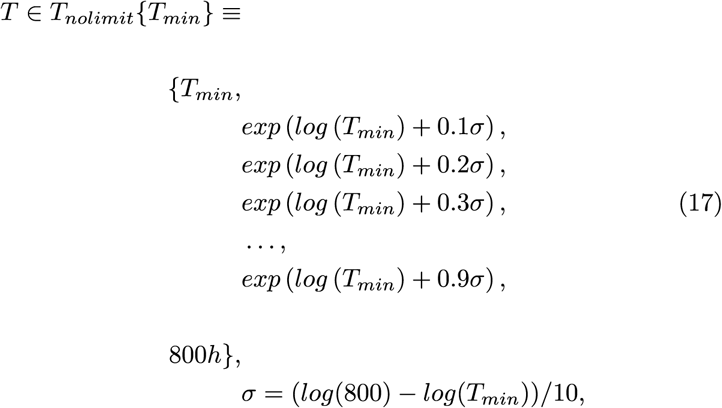

and *T*_*min*_ = 256*h*. The set of occupation probabilities *p* defined in equation (16) will be used in all the FTS procedures throughout this work.

The Spearman’s Rank Correlation Coefficient Optimization method (SRCCO, see section 5.7) was implemented to collapse the *R* (*T* |*p*)∼ *p* curves and obtain the critical SPR scaling relation 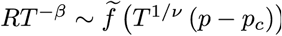, where *f* is the scaling function, *β* and *ν* are the scaling critical exponents, and *p*_*c*_ is the critical occupation probability. It could be observed that the *R* (*T*| *p*) ∼ *p* curves with *T* = 713.8h and *T* = 800h in Fig. S2a are almost overlapping on top of each other. The greater the time scale, the greater the extent of the overlapping, which conflicts with the FTS. It has caused the failure of the FTS analysis shown in Fig. S2b. Fig. S2b shows the result of collapsing the *R (T*|*p)* ∼ *p* curves through the SRCCO method. It could be deduced that SRCCO would almost fail to collapse the *R*_*i,j*_ points together for *T*_*i*_ ∼ 800h. This is because it can be inferred from the *R* (*t*|*p*_*j*_) ∼ *t* curves in Fig. 1 that *R (t*|*p*_*j*_) approaches a constant as *t* approaches infinity, defined as *R(∞*|*p*_*j*_) ≡ *Const* (see Supplementary Material S3), and thus causing the total overlapping of the *R (T*|*p)* ∼ *p* curves. It is further explained in Supplementary Material S3 why this leads towards the conclusion that SRCCO will almost fail to collapse the *R*_*i,j*_ points for *T*_*i*_ ∼ 800h and how to determine an upper limit to the finite system time scale *T* for guaranteeing sufficient collapse quality. This upper limit is denoted as *T*_*max*_, and called the ‘Max Permitted Scale’ (MPS). The optimal time scale set for collapsing curves is denoted as *T*_*aptimaz*_*(T*_*min*_, *T*_*max*_). The set of the critical values derived with *T*_*aptimat*_*(T*_*min*_, *T*_*max*_) is denoted as *ϕ*_*optimal*_*(T*_*min*_, *T*_*max*_*)-* For an illustration of these notations, see Supplementary Material S3.

### 3.3 Negative correlation between the invasiveness and the transition point p_c

To test its measurement capability, the FTS analysis was implemented on virtual cancer cell populations of three parameter sets. The invasiveness of these virtual cancer cell populations can be compared with each other by intuition.

Fig. 2a shows the *R (T*|*p)* ∼ *p* curves, an intuitive sense of SRCCO collapse quality could be obtained by comparing these curves with the collapsed ones in Fig. 2b. It could be observed that the fanning out curves in Fig. 2a have almost been collapsed into a single curve. The critical exponents thus derived are *p*_*c*_ = 0.3408±3*E* − 4,*ν* = 4.236 ± 9*E*−3, *β* = 1.917±1*E* − 3, respectively.

**Fig. 2.**
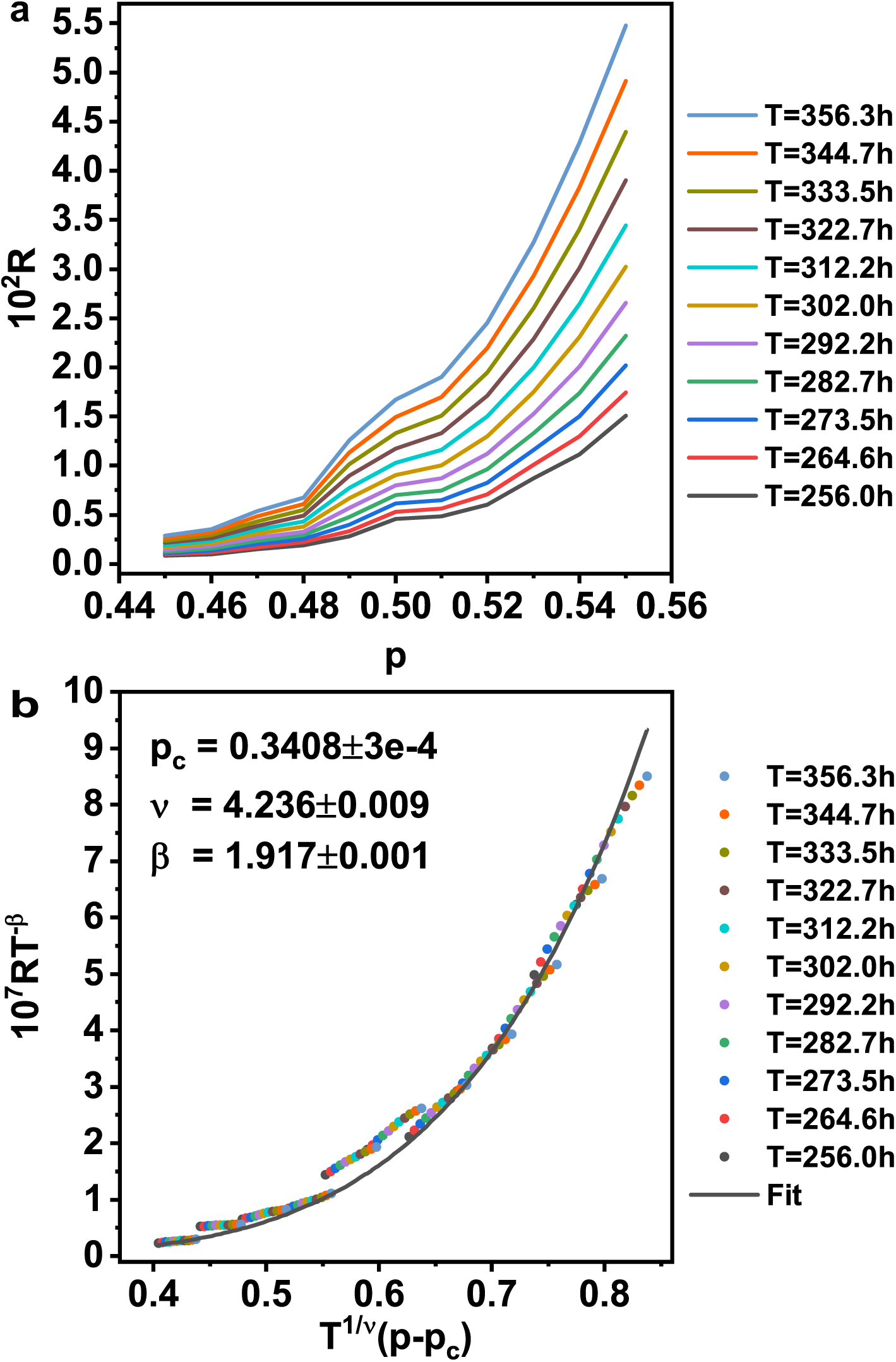
The FTS result of the model fitted to the raw data. Figure a shows the relationship between *p* and *R, T*_*i*_ ∈ *T*_*optimal*_ (256,356.3). Figure b is the FTS result with the critical values *p*_*c*_,*ν*, *β* generated by SRCCO fed with the SPR data *R*_*ij*_, *i* ∈ *p, T*_*i*_ ∈ *T*_*optimal*_ (256,356.3). The discrete points correspond to 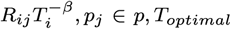 (256,356.3). The grey curve is the fit for those discrete data.

Fig. S4 in Supplementary Material is the second FTS subject, with the paracrine signaling paths removed. Specifically, it was set that *µ*_1_ = 0. By intuition, this virtual population should be less invasive than the original one. The critical exponents thus derived are *p*_*c*_ = 0.3675 ± 3*e*−4*, ν* = 3.886 ± 8*e* − 3, *β* = 1.7472 ± 3*e* − 4, respectively.

After a few more modifications, a third FTS phenotype is produced, as now the paracrine signaling pathways have been removed and the death probability has been tenfold increased. Specifically, *µ*_1_ = 0, *ϵ* = 10*ϵ*_*original*._ By intuition, this virtual population should be less invasive than the both ones above. The critical exponents thus derived are *p*_*c*_ = 0.4454 ± 2*e* − 4, *ν* = 4.871 ± 0.002, *β* = 2.0470±6*e*−4, respectively. Shown in Fig. S5 in Supplementary Material is a further illustration.

From such observed negative correlation between *p*_*c*_ and the invasiveness, it was inferred that the framework designed here could measure the invasiveness into a 2D-percolation cluster. From that *p*_*c*_ has gone from 0.34 to 0.36 and then to 0.44, it could be inferred that as invasiveness weakens, a cell population needs greater occupation probability to grow. Therefore, this suggests that inferior adaptivity to dense ECM goes hand in hand with weaker invasiveness. Given the abovementioned considerations, the conclusion was drawn that *p*_*c*_ might be promising to be further exploited for invasiveness measurement.

A hypothesis on the underlying mechanism is that the cell population gets more insensitive to the overall site configuration due to a lack of ubiquitous paracrine signals connecting the individual cells. Without paracrine signaling, populations can only sense permeability change with mechanical interaction, while those with the signaling have a long-range alternative. For populations capable of paracrine signaling, the paracrine signals get hindered by site configurations as well as the cells do. And these two types of geometrical obstruction may take effect in a synergetic way. This may have caused the higher sensitivity to occupation probability. The further details are shown in the Discussion.

In a nutshell, the results above suggest that the critical behaviors of the populations could get significantly affected by the intensity of the paracrine signaling. Note the paracrine signaling paths represent the inter-cell operation that leads to the emerging Allee effect (Korolev et al, 2014), so it can also be concluded that the critical behaviors could be significantly affected by the intensity of the Allee effect.

## 4 Discussion

It was mentioned in the Introduction that the emergence models of the Allee effect have a huge parameter space and multiple behavioral patterns. Most of those patterns have not been verified by experiment. A better understanding of the underlying principles was hindered by the complexity of the interactions and the limited control of the parameters in the existing experiment frameworks.

Accurate correspondence between theory and experiment could be achieved by choosing site configuration as the control parameter, as it will become a highly controllable geometric condition in the future (Diao et al, 2019). This is important because it will be convenient for the researchers to carry out accurate control, so that they could arrange in experiments the same initial conditions proposed in theories for parameter inference. In this way, a virtuous cycle could hopefully be set in motion between theory and experiment.

Here the FTS analysis was carried out on the Growth-Migration processes of cancer cell populations in 2D percolation cluster. The critical exponents were derived for further analysis.

### 4.1 The necessity of the MPS *T*_*max*_

The necessity of *T*_*max*_ stems from the rationale that the collapse procedure should not be carried out for a collection of curves almost not possible to be collapsed, e.g., the curves in Fig. S2a in Supplementary Material.

Inference 1 in Supplementary Material S3 suggests that when *T* → ∞, *R (T*|*p)* → *Const*. The premise of Inference 1, i.e., Supplementary Material Equation (S.1), was obtained through observing the results in Fig. S2 in Supplementary Material. Nevertheless, it could also be deduced analytically. It might as well be assumed that (Assumption 8) The cell death propensity, *ϵ*, could be assumed to be sufficiently low so that the population will not go extinct.

Such an assumption is based on the fact that for both the original model and the variations, the mitosis propensity *γ*_0_ = (2.7 ± 0.09) *e* − 6 *s*^−1^ and *γ*_1_ = (3.3 ± 0.04)*e* − 6 *s*^−1^ are way greater than the death propensities *ϵ* = (7.4 ± 4.9)*e* − 8 *s*^−1^ or 10*ϵ*= (7.4 ± 4.9)*e* − 7 *s*^−1^. Therefore, as *t*→ ∞, ∀*j, k*,

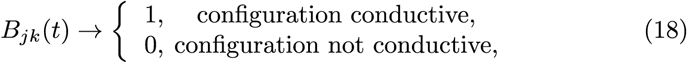

thus,as *t* → ∞, ∀*j, F(t*|*p*_*j*_) →Const, and so, *R(t*|*p*_*j*_) ≡ < *F* > *(t*|*p*_*j*_) → Const.

SRCCO is barely able to collapse the *R(T*|*p*)_*T*∼800_ curves with assumption 8 satisfied.

In conclusion, the concept of the Max Permitted Scale *T*_*max*_ was introduced to assure the exclusion of the curves that may render the SRCCO procedure nearly impossible.

### 4.2 The meaning of the critical exponent

After The definitions of the critical exponents depend on the actual behaviors they depict. Following are the critical behaviors under consideration.

Supposing a quantity *A*_*T*_ (*p*) diverges in the manner *A*_*T*_ (*p*) ∼ |*P* – *p*_*c*_|^−*ζ*^ in the critical region of an infinite system *(T* → ∞, *p* → *p*_*c*_), thereby the definition of *ζ, p*_*c*_ was obtained.

The characteristic return time (Sorge, 2018) is denoted as *ξ* (*p*). In one-dimensional temporal space, *ξ* (*p*) is also the temporal coherence scale of the infinite system. And so *ξ* (*p*) behaves in the way *ξ* (*p*) ∼ | *P* − *p*_*c*_|^−*ν*^, *(p* → *p*_*c*_), thereby the definition of ν was obtained.

For further details, see section 5.6.

### 4.3 On the comparison among the three virtual cancer populations with varied parameter sets

The set of fitted parameters here could serve as a prototype of the spatiotemporal in-vitro model, as it could well fit the mean temporal trajectories of the total cell number. Tuning certain parameters may help to observe the critical behaviors of populations with varied invasiveness. In other words, this model might be a small step towards simulating virtual in-vitro culture assays under various perturbations.

The comparison among the three model variations mentioned in S ection 3.3 inferred that *p*_*c*_ may serve as a quantitative indicator of the invasiveness This interpretation could be indirectly supported by the work of (Caicedo-Carvajal et al, 2011) on the lymphatic cancer cell culture in 3D lattices. They had found that both 2D and 3D geometry is not relevant to the growth of the population when there had been no stromal cells. Nevertheless, growth was greatly enhanced after introducing the stromal cells into the 3D lattices but not for the 2D ones. They have postulated that the reason might be that certain factors secreted by the stromal cells had activated the genetic switch of the cancer cells. Consequently, the cancer cells had become more likely to interact with the stromal cells. Note that the total area of both 2D and 3D was designed to be the same. Here, it could be guessed that, with the same total area, a population in a 3D lattice is more compact, leading to more efficient signal dissemination across the whole region so that the perk brought about by the stromal cells stands out in 3D lattices rather than 2D ones. That is why the work may support the intuition that ‘paracrine signaling promotes the invasiveness’.

#### 4.3.1 Framework with promising scalability

Critical exponents with low uncertainty were obtained by the FTS method. Subsequently, the mathematical model must be improved for greater robustness of parameter inference. Only structurally identifiable parameters can make it possible for theoretical computation to take the place of large-scale ensemble assays. Then, critical exponents could be obtained by feeding the computation results into the FTS software SJLY(SJLY Shao-Jiang-Liang-Yang) designed in this work. SJLY is freely available at http://www.urdme.org.

Therefore, it is high time a more detailed model be developed. Thanks to inspiration by (Engblom, 2019), a primitive framework with promising scalability is proposed conceptually in this section, which may be realized in the future by developers interested.

Coupling models could achieve scalability at multiple time scales in continuous time. And it is proposed here that this could be achieved by nesting the Next Subvolume Method (NSM) simulations (Drawert et al, 2012) at varied time scales. Engblom (2019) has managed to couple a PDE model with an NSM model by nesting, but no framework nesting NSM models was mentioned. The concept of NSM nesting is demonstrated in Fig. 3. Shown in Fig. 3 are three NSM procedures at varied time scales. The time point notations in Fig. 3 were simplified by the definition

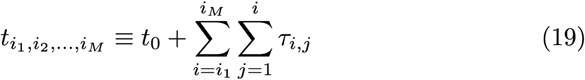

**Fig. 3.**
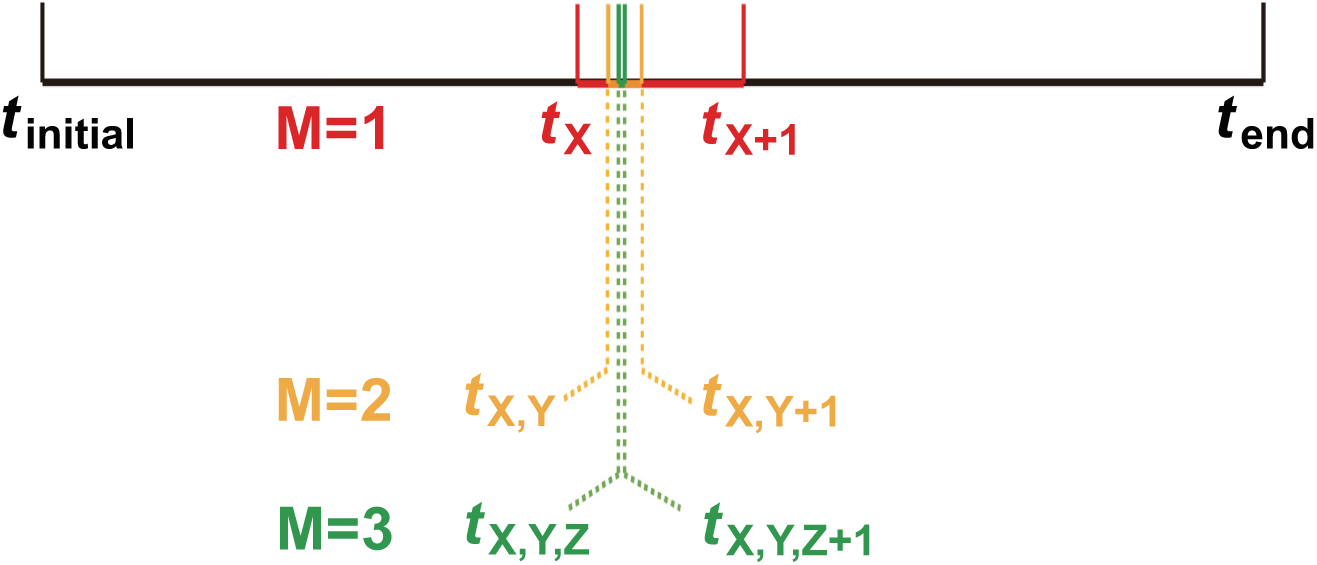
A scheme of three NSM models nesting together, *N* = {*N*_1_, *N*_2_, *N*_3_}. Black segment represents the total simulation time. The red, yellow, green segments represent the intervals sampled in the *X*th iteration of *N*_1_, in the *Y*th iteration of *N*_2_, in the *Z*th iteration of *N*_3_, respectively. The time point notations coresponding to the ends of the segments denote the ends of the intervals.

Let us define the order of an NSM model as its ranking among a set of NSM models according to the time scales, and let us, e.g., determine that a lower time scale leads to a lower order. Consider a set of NSM models:

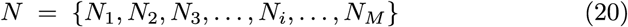

where N denotes the set itself, *N*_*i*_ denotes the *i*th order NSM model, and M denotes the highest order among the elements. *i*_*n*_ (*n* = l, 2, …, *M)* denotes the accumulated numbers of iterations of *N*_*n*_ done during a simulation. *τ*_*i,j*_ denotes the interval sampled during the *j*th iteration in the *i*th order NSM procedure. *N*_*n*_ (*n* = 1, 2, …, *M*) could run on itself regardless of each other, until it prepares to go beyond the interval sampled by *N*_*n*+1_, i.e., until it prepares to go to the regions on the time axis where the information of its ‘adiabatic’ environment has not yet been decided by *N*_*n*+1_. This waiting scenario may happen if the NSM models require information from each other, thus synchronization may be required when an iteration is over.

One may categorize the time scales into four types: cellular scale, macro-molecular scale, micro-molecular scale, and quantum scale.

This research has two main limitations. The identifiability of the parameters of the model here is limited by the coarse grain phenomenological approximations of the signaling mechanism, and inadequate depiction of the forces exerted on the cells. A coarse-grain pseudo-particle method was used to describe the signaling pathway. Secretion of mitogenic signaling molecules within a period was represented by the release of a signal package within the same period. This is a quite rough approximation. On mechanics, intercell mechanical interactions was simply neglected. Under the circumstances where cells collide into each other within narrow space, such an approximation is flawed.

In conclusion, this work serves as an example of how to use FTS to characterize the spatiotemporal stochastic processes of a population in a quantitative way. The SPR transition point defined according to the FTS theory, *p*_*c*_, was found to be negatively correlated with the invasiveness of designated virtual cancer cell populations. Therefore, the FTS algorithm may be useful in determining invasiveness. Through the simplistic phenomenological paracrine model, inter-cell cooperation and mutual mitogenic boosting are enabled, causing the Allee effect to occur. Such a method could be applied to other circumstances as an example of the quantitatively falsifiable emerging theory. To generate samples of percolation cluster, additional details and large-scale computations are needed. However, controllable geometric properties can serve as the control parameter, and the SPR could serve as the observable for its accessibility and precision. Through these settings, the degree of uncertainty of observation can be greatly reduced, and the repeatability of the experiment can be improved, thus strengthening the foundation for parameter inference and discovery confirmation.

## 5 Material and methods

An agent-based URDME model (Drawert et al, 2012) set in a 2D culture well was fitted with the experimental data of the in-vitro culture assays of (Johnson et al, 2019). All codes of the model were included in the PAllee software package provided here. With this model, the evolution of the spatial distribution of the cells can be obtained as well as the temporal trajectories of the total cell numbers. The data for parameter fitting was the mean temporal trajectories of the total cell number obtained by (Johnson et al, 2019). The fitting algorithm relied on the global searching by a genetic algorithm method. The precision of the fitting algorithm was measured by the uncertainty of repeated searching. The discoveries in this work are based on the repeated computation and statistics of the model.

### 5.1 Algorithm of computation

URDME (Drawert et al, 2012) addresses the stochastic Finite Element Method. It has achieved computation of stochastic processes while inheriting the meshing algorithm of the Finite Element Method (FEM). It uses the dual mesh of the triangular mesh of the FEM software COMSOL. The relationship between the dual and triangular meshes is shown in Fig. 4. A dual mesh is obtained by connecting the geometric centers and the midpoints of the edges of each triangle in the triangular mesh. Each polygonal, e.g., the red-edged one in Fig. 4, is called a ‘subvolume’. URDME represents the position of each subvolume with the original triangular node in the center of the subvolume.

**Fig. 4.**
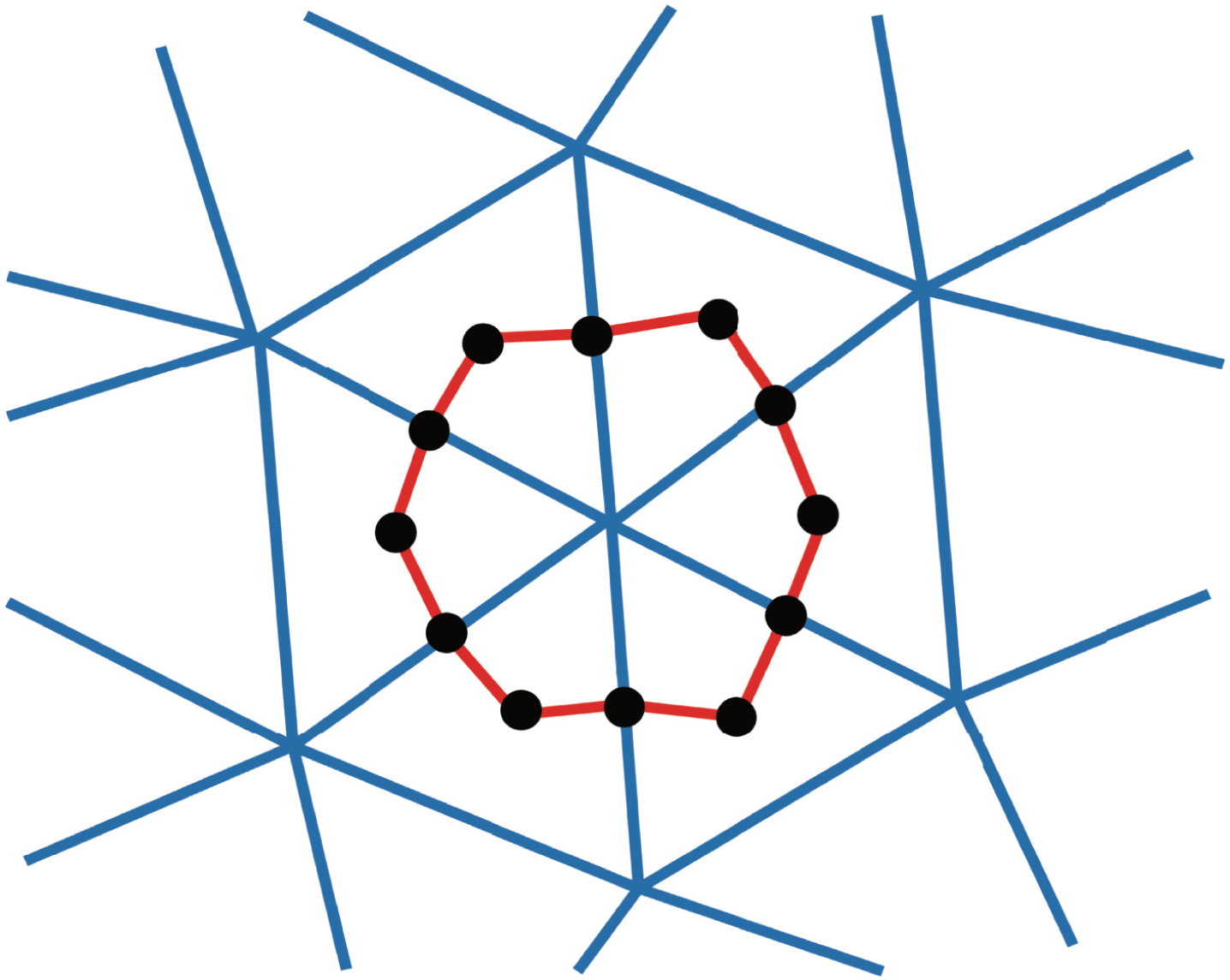
The figure shows how a triangular mesh is converted into its dual mesh. The dual mesh is obtained by connecting the geometric centers and the midpoints of the edges of each triangle in the triangular mesh. The geometric centers and the midpoints are represented by black dots. Edges of the dual mesh are represented by red line segments. For clear demonstration, only the dots and edges corresponding to one of the subvolumes are shown, and the rest are all hidden.

URDME is a type of agent-based method, also known as an ‘Next Subvolume Method’ (NSM) (Drawert et al, 2012). NSM got its name because both the start and end of the next border-crossing event in equation (11) are picked according to the Poisson rate parameters 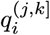, by the ‘inversion generation’ sampling method described in (Gillespie, 1976). In this ‘next subvolume’ ‘inversion generation’ sampling procedure, the next subvolume for departure is first determined by drawing lots among the qualified candidates, and then comes the determination of the next subvolume for destination in a similar way. By the way, the destination subvolume must share the same edge with the determined departure subvolume.

### 5.2 Geometry settings

To compute the evolution of the total cell number, geometrical abstraction is made in COMSOL for one of the wells within the 96-well culture plate (Trueline Saint-Anne-de-Bellevue, Quebec, Canada), as shown in.Fig. 5. The mesh in the figure was generated automatically by COMSOL. A relatively finer (50 *µm*) mesh was used for the 1mm*1mm square region in the center to save computational resources. This is plausible in that it was suggested by the green fluorescence photos of the cell culture assays in (Johnson et al, 2019), that in a culture well, most of the cells had moved within a 1mm*1mm range around the seeding point during the first 328 hours. It is this region that is of interest.

Following are the concepts of ‘Typical Size of Interest’ (TSOI) and ‘Typical Subvolume’.

**Fig. 5.**
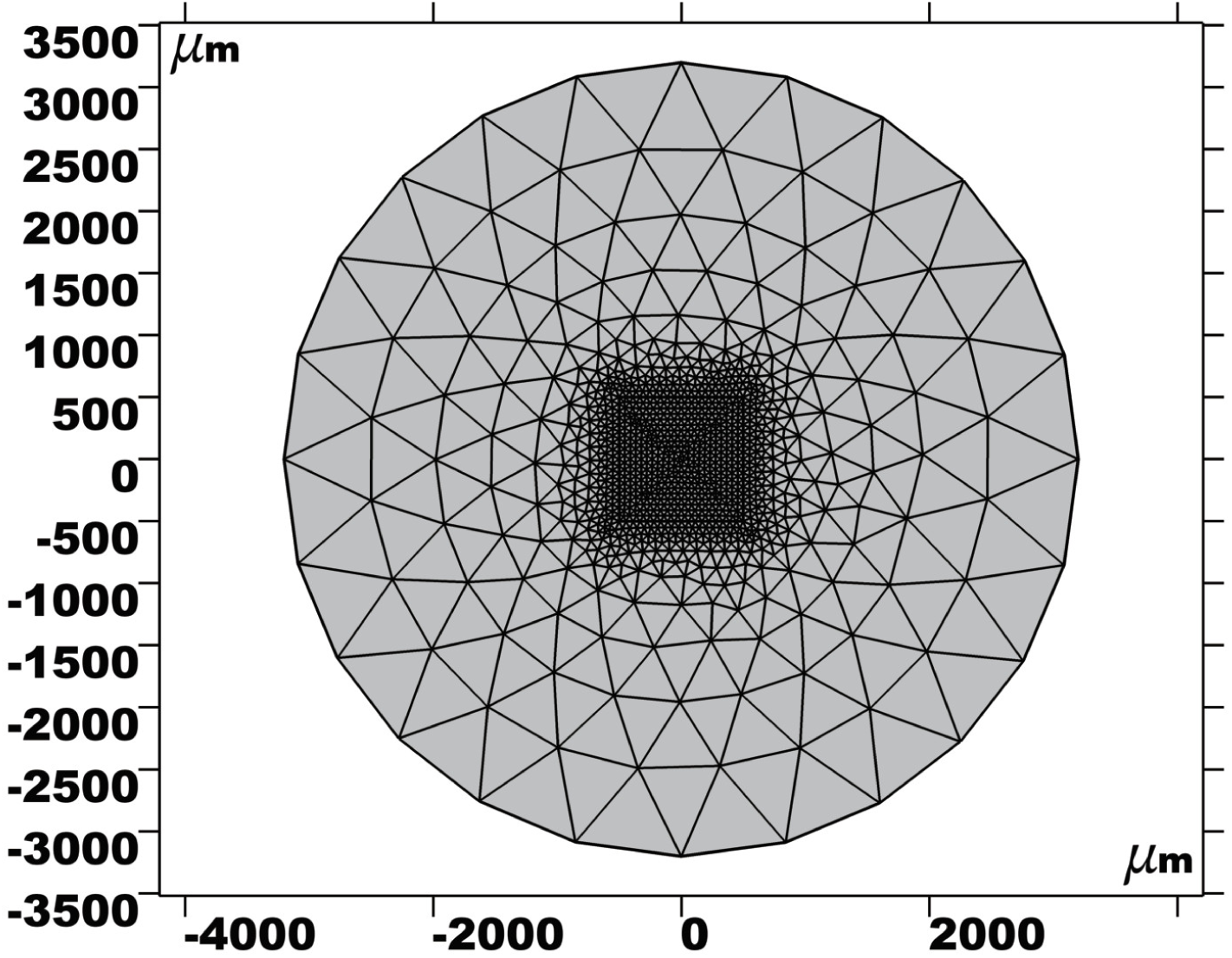
Geometrical abstraction made in COMSOL for one of the wells within the 96-well culture plate (Trueline Saint-Anne-de-Bellevue, Quebec, Canada), The mesh was generated automatically by COMSOL. A relatively finer (50 *µm)* mesh was used for the 1mm*1mm square region in the center.

The relationship between them is shown in Fig. 6. Hard circular disks were used to model the cells (Fig. 6d). It could be inferred from (Cadart et al, 2018; Cleris et al, 2019) that the effective diameter of the hard disk models of S_1_ or S_2_ should be set as 20 *µm*. For convenience, this effective diameter is called the ‘Typical Size of Interest’ (TSOI).

**Fig. 6.**
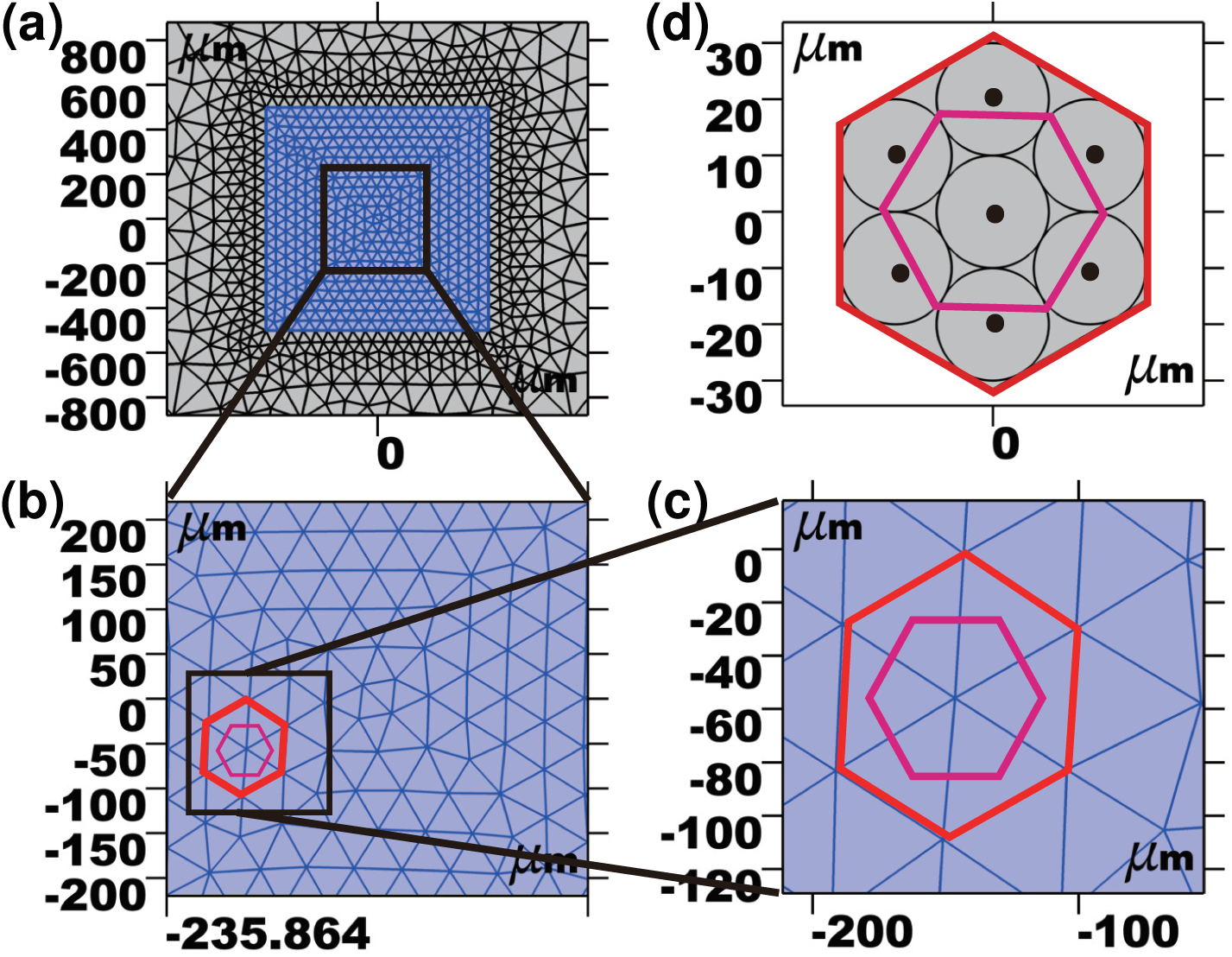
The blue region in 6a represents a 1mm*1mm square. It is treated as the region of interest. The black square is a region randomly selected for zooming in. 6b shows the enlarged area of the black square in 6a. The magenta polygon within the red polygon represents a typical subvolume. It could be inferred that it is almost a regular hexagon. The red polygon represents the nearest edges of the triangular mesh around that typical subvolume. The black square in 6b serves in the same way as that in 6a. 6c is an enlarged version of the area chosen by the black square in 6b. 6d shows the relationship between the TSOI and an ideal typical subvolume. The magenta regular hexagon represents an ideal typical subvolume, and the red regular hexagon represents the nearest edges of the triangular mesh around it. These edges also form a regular hexagon. The circles represent the hard disk models with the TSOI as their radii. The black dots represent the center of the hard disks.

The subvolumes within the 1mm*1mm region of interest resemble each other, almost identically equal. Therefore, they were called the ‘Typical Subvolumes’. The size of the typical subvolumes was designed by setting the size of the triangular mesh. Shown in Fig. 6d is the ideal relationship between the TSOI hard disks and an ideal typical subvolume. This ideal typical subvolume is a regular hexagon. As shown in Fig. 6d, the triangular mesh edges (red) encircle an ideal typical subvolume and form a regular hexagon. The size of this hexagon is designed in such a way that exactly seven TSOI hard disks will make it stuffed. The size of the dual triangular mesh of the ideal typical subvolume is called the ‘Ideal Typical Triangular Mesh Size’ (ITTMS).

### 5.3 Details of the model

Detail 1 Inspired by the observations in (Boucrot and Kirchhausen, 2008; Cadart et al, 2018), it was assumed that the effective area of a daughter cell in 2D geometry maintains half of that of its mother in concomitance with the occurrence of the event in equation (5.2/2), and then grows into a fully grown cell in concomitance with the occurrence of the event in equation (5.2). This could be expressed with symbols that the area of S3 equals to 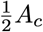. So the total cell area within a subvolume is (*X*_1_*A*_*c*_ + *X*_2_*A*_*c*_ + 0.5*X*_3_*A*_*c*_).

Detail 2 A restriction on proliferation. The proliferation will be prohibited when a subvolume gets overcrowded. The crowdedness is represented by the proportion of total cell area (*X*_1_*A*_*c*_ + *X*_2_*A*_*c*_ + 0.5*X*_3_*A*_*c*_) to the subvolume area *V*. When this ratio gets less than *ρ*_1_, the propensity coefficient, *γ*_0_ and *γ*_1_ will remain constant. And when it surpasses *ρ*_1_, it is determined that the subvolume undergoes overcrowding, and then the propensity coefficient *γ*_0_ and *γ*_1_ will be set to zero. Finally, because *γ*_1_ is for the boosted cells, naturally *γ*_1_ > *γ*_0_ The restriction in the form of equations is as follows:

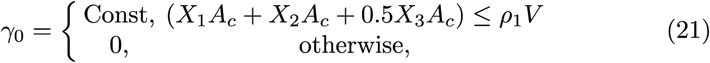

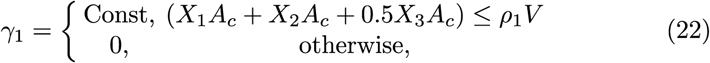

Detail 3 The mitogenic signaling was modeled as the signal packages. In this way, the behaviors of only a single species *S*_4_ can be used as an equivalent of both the diffusion and cellular interactions of all the known and unknown mitogenic signaling molecules. Such description could phenomenologically and yet quantitatively depict the mitogenic signaling paths. Interpreting the signaling molecules as signal packages is a natural choice. This idea had been inspired by how Jin et al (2020) converted the molecule density field within a subvolume into a local mean-field. The intensity of a mean-field within a certain subvolume was a function of the cell number within the same subvolume. The relaxation time of the mean-field was tuned to be consistent with that of the cell number. This coarse-graining thinking was followed here. Secretion of mitogenic signaling molecules within a period was represented by the release of a signal package within the same period. Abstract as such depiction is, it serves as a straightforward way to describe signaling pathways of in-vitro cell cultures.

Detail 4 A restriction on migration. When the initial seeding number is greater than or equals 10, it is possible that overcrowding occurs within some subvolumes during the whole simulation. This is solved by putting a restriction on the propensity coefficients 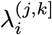, of the border-crossing events. If the initial cell number is less than 10, then the Poisson rate parameters will be computed as usual. If it is greater than or equal to 10, then it should be checked whether this subvolume belongs to the collection of the subvolumes undergoing overcrowding, i.e., 𝕍_*OC*_

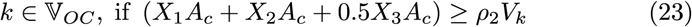

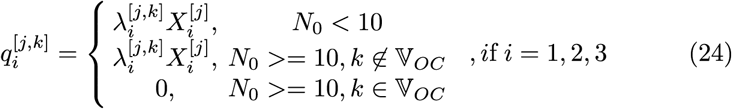

Detail 5 For diffusion of the signal packages, it was assumed that there is no collision between the packages. Coarse-graining and rough as it is, this is still the simplest approach:

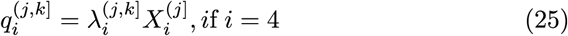

Detail 6 It was assumed that the border-crossing propensity coefficients are the same for *S*_1_ and *S*_2_ to simplify the model, i.e.,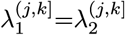.

### 5.4 Experimental data for parameter fitting

The agent-based model set in a 2D culture well here was fitted to the in-vitro culture data by (Johnson et al, 2019). The Growth-Migration evolution of a cancer cell population within a culture well could be depicted with this model.

Following is a detailed illustration of those data.

The data of the temporal evolution of three assays with seeding numbers 2/4/10, respectively (Johnson et al, 2019), was considered here. They were illustrated in figure 9 in the original work. Their experiment was carried out in planar space. Initially, the cancer cells were seeded in the circular culture wells, and the media were then renewed regularly. The weak Allee effect was confirmed by fitting the stochastic models to the data of LSDC assays.

A substrain of the BT474 breast cancer cells was designed to consistently express enhanced green fluorescent proteins, which carry the location signal of the nuclei. The cells were cultured on a pre-coated media and then seeded with initial seeding numbers at the precision of a single cell. The LSDCs were then grown in the media for seven days. Subsequently, the media were refreshed every 2 to 3 days in the following two weeks. Pictures were sampled every 4 hours during these two weeks. The actual initial cell numbers (2/4/10) were confirmed by eyes analyzing four-fold magnified pictures, and the temporal evolution trajectories of the cell numbers were also binned in this way.

### 5.5 Method of parameter fitting

An objective function was designed to be optimized by the Matlab Genetic Algorithm toolbox ‘ga’.

To estimate the distance between the true values and the candidates, a first order moment closure method (Fisher, 1922; Froehlich et al, 2016; Raue et al, 2009, 2013) was implemented in the objective function design.

Following is the objective function:

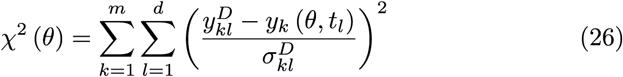

where the label “D” represents “data”, i.e., the experimental data, 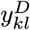 denotes the data point of the *k*th observable measured at a time point *t*_*l*_, 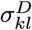 denotes the corresponding measurement error, *y*_*k*_ (*θ, t*_*l*_) denotes the data point of the *k*th observable computed for time point *t*_*l*_ by the model with parameter *θ*. The true values of the parameters can be estimated in the following way, i.e., choosing the parameter 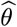 that minimizes the objective function.

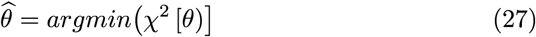

For a normal observable noise 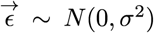, the objective function *χ*^2^ *(θ)* corresponds to the maximum likelihood estimate (MLE):

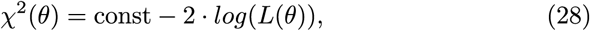

where *L(θ)* is the likelihood function in the MLE theory. *χ*^2^ will be used to represent likelihood in the rest of the article.

The fitting was carried out in a 2D virtual culture well consistent with the in-vitro culture assays. The boundary condition is a no-flux condition corresponding to the actual boundary of the wells in the assays. The signaling packages are described by discrete agents, and the propensity coefficients of the Poisson events that they take part in have all been directly fitted to the experimental data. This has guaranteed a parameter range with biological significance. Tuning the parameters within such a range could lead to more realistic quantitative computational results or qualitative observations.

The parameters for fitting and depicting the Growth-Migration within a circular culture well are then used to describe the Growth-Migration within a 2D percolation cluster.

With the estimated parameters, the computation could be taken as a virtual experiment. The price for adding two spatial dimensions to the temporal dimension is more parameters–Johnson et al (2019) have used only four parameters to model the stochastic equations for the total cell number evolution. However, the uncertainty was contained within a relatively small range through large-scale repeated simulation batches for fitting the experimental ensemble mean. Johnson et al (2019) have obtained the mean temporal trajectories of the total cell number with different seeding numbers by analyzing a huge number of repeated assays. And then, they used the moment closure method and obtained parameters with sufficiently low uncertainty by fitting the stochastic models to the experimental data.

The ‘ga’ toolbox was used to optimize *χ*^2^*(θ)*. Although ‘ga’ has already been a global optimization method itself, three repeated runs were carried out to analyze the uncertainty of the Max Likelihood Estimation (MLE) 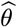. The fitted MLE and the corresponding uncertainty can be seen in Table. S1 in Supplementary Material S1

### 5.6 Design and generation of random percolation cluster

Any FEM nodes at the boundary need a no-flux boundary condition employed to establish the FEM mesh of a 2D percolation lattice. This was achieved by tuning the border-crossing propensity coefficients 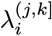 in the software ‘SJLY’ developed in this work. Specifically, the no-flux condition was set by prohibiting any agents from moving into the FEM nodes on the boundary. The rationale behind, as illustrated in Fig. S6 in Supplementary Material, is that nonzero cell numbers on the boundary lines and boundary corners will enable cells to move between unconnected sites through their common corner nodes, thus leading to great errors. Of course, a price for this method is that the subvolumes centered by those nodes are rendered inaccessible. However, unfortunately, in each percolation site, it is desired that at least one subvolume centered by a non-boundary node should exist. Therefore, given the TSOI of the subvolume, the side length of each site was designed to be twice as that of the TSOI, i.e., 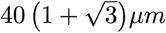.

With the software SJLY, percolation lattices were generated by Monte Carlo rules. For a given occupation probability *p*, whether each site is occupied is determined by a new temporarily generated random number *U*(0, 1).

### 5.7 Finite time scaling analysis

As shown in Fig. 7, Sorge (2018) has proposed the FTS theory for general stochastic processes. The theory claims that the return of the stochastic processes can be described by one-dimensional percolation on the temporal axis. By ‘return’ he meant a return event, i.e., the state returning to the initial state. By such mapping, vacancy emerges between the returns, cutting the continuous temporal axis into discrete clusters. The size of each cluster is equivalent to the period between two returns, which is named the return time, denoted as *τ*.

**Fig. 7.**
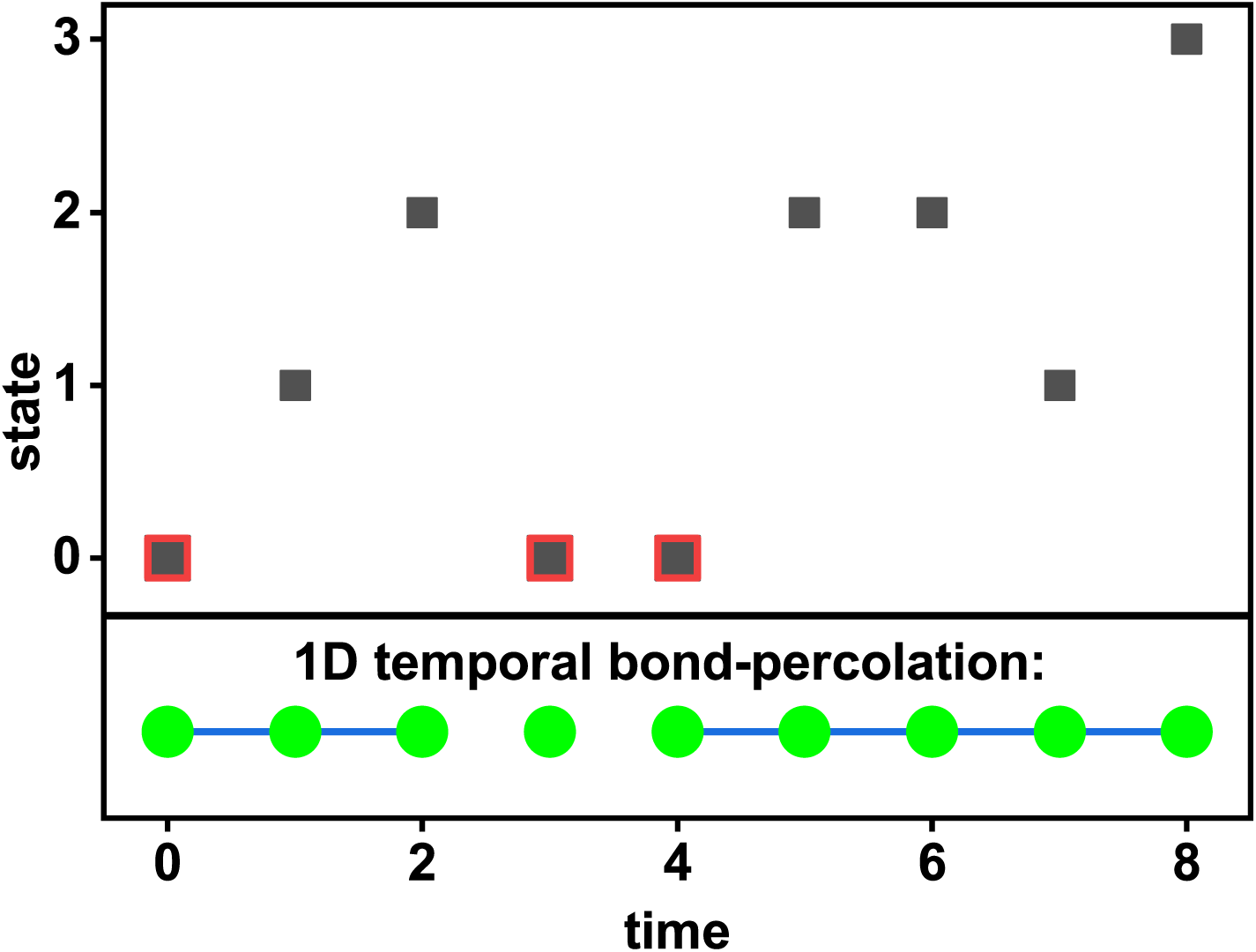
Andrea Sorge mapped the return dynamics (the black part) onto a one-dimensional bond percolation model (the green part in the bottom). Depicted in the figure is a sample of the first eight time steps of the random walk on the half-line. The highlighted squares correspond to the returns at time points 0, 3, 4, respectively. In the bond percolation model at the bottom, each site corresponds to a time point. A bond emerges along the step during which no return occurs.

One-dimensional percolation was introduced by (Sorge, 2018). Bond breaking is supposed to be triggered by the onset of returns. Cancer populations could fail to grow up to a threshold and wither afterward. Return can be used to model such recessive processes of the Growth-Migration of cancer populations. Therefore, this theory may provide quantitative and verifiable analysis of the Growth-Migration dynamics.

The distribution of the return time could serve as the basic quantity of one-dimensional percolation. As how the scaling theory links cluster sizes with the percolation transition critical exponents, (Sorge, 2018)’s theory of the external model links the avalanche size distribution with the critical exponents of self-organized spatiotemporal dynamics. Sorge (2018) has linked temporal percolation transition with the avalanche size distribution following this rationale.

Based on the stability theory of the Markov chain and the critical transition theory of (Maslov, 1996; Stauffer, 1979), Sorge (2018) has hypothesized a phase transition of a Markov chain with a parameter *p* and supposed that the transition featured a return time distribution with exponential behavior as:

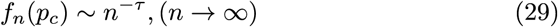

where *τ* is the Fisher exponent, *n* the return time and *p*_*c*_ the critical transition point. The scaling hypothesis is that the characteristic return time is *ξ(p)*, and that 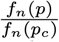 is only a function of 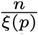. The characteristic return time *ξ (p)* is equal to the temporal coherence scale of an infinite system, as it is a onedimensional space that is under consideration. So *ξ* (*p*) is supposed to behave as

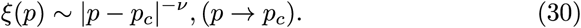

Finite time simulation cannot probe the critical transition directly because the transition only occurs in an infinite system. For a finite time system, the regular remedy for finite size scaling (FSS) analysis was borrowed here for addressing one-dimensional temporal percolation. The finite time scaling (FTS) analysis infers the critical point *p*_*c*_ and the critical exponents by numerical algorithms.

*T* denotes the time range of a dynamic system, i.e., the total number of time steps in the computation. *A*_*T*_*(P)* denotes a quantity converging as |*p* − *p*_*c*_|^−*ζ*^ within the critical region *(T* → ∞,*p* → *p*_*c*_) of an infinite system. The FSS ansatz could be interpreted according to (Binder and Heermann, 2010; Fisher and Barber, 1972; Newman and Barkema, 1999) as

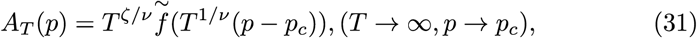

where 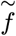 is a dimensionless scaling function, *ν* is the critical exponent for the characteristic temporal coherence scale *ξ* (*p*) in an infinite system. The finite time effect is supposed to be controlled by the scaling function.

Suppose a series of data were obtained, denoted as 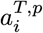. *i* is for the *i*th computational result, *T* the system size, and *p* the parameter. *a*_*T,p*_ is used to represent the mean of each computational result. The numerical computation data should collapse onto one master curve 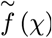, with 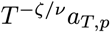 as the vertical axis, and 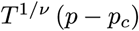 the horizontal axis. According to (Binder and Heermann, 2010; Newman and Barkema, 1999), the hypothesis hold for T → ∞. Finite time might lead to errors, and customized adjustments are needed to achieve the collapse.

For convenience, *ξ/ ν* is simplified as *β*. The collection of the critical values is defined and denoted as Φ ≡ {*p*_*c*_, *ν, β*}.

### 5.8 Curve collapsing algorithm SRCCO

According to the FTS method proposed by (Sorge, 2018), the step after obtaining the *R* (*T*|*p*) ∼ *p* curves is collapsing the curves into a master curve. For more about the rationale behind the collapse procedure, see (Binder and Heermann, 2019; Newman and Barkema, 1999). The method implemented here includes the discrete points on all these curves in a single set and then computing the Spearman’s Rank Correlation Coefficient (Spearman, 1904) of this data set. See also the statistics dictionary by Springer press (**?**) for a definition of the Spearman Correlation. **?** have implemented FSS analysis when studying Growth-Migration of cancer populations exhibiting Allee effects within 2D percolation cluster. They used Spearman’s Rank Correlation Coefficient to assess the extent to which the candidate critical exponents collapse the raw curves, and then feed it as an objective function into an optimization algorithm. As a result, they found that there existed a transition point of the SPR with the occupation probability as the changing parameter. Such a curve collapsing algorithm is categorized here as Spearman’s Rank Correlation Coefficient Optimization (SRCCO) algorithm. A software implementing SRCCO, SJLY, is launched here. A user can feed the set of the temporal trajectories of the Boolean values *B* = {*B*_*jk*_(*t*),*j* = 1,2,3…; *k* = 1,2,3, …; *t ∈ T*_*sim*_} and then directly get the critical values as output, where *Tsim* is a user-customized set of the time points for result sampling.

## Authors Contributions

RY and YS conceived and performed the study. RY, YS and CJ wrote the manuscript.

## Availability of Data and Materials

The SPR trajectories and the percolation configurations can be found at https://pan.baidu.com/s/lmksIOb2EMcAG9p7j62MksA?pwd=y4xh

## Code Availability

The code of PAllee and SJLY can be found at https://github.com/RenlongYang

