## Supplementary Information for "Invasiveness of Cancer Populations in a Two-dimensional Percolation cluster: a Stochastic Mathematical Approach"

#### S1 Fitted parameters of the PAllee model

See Table. S1.

#### S2 An illustration of the percolation assay settings

Shown in Fig. S1 is an example of the evolution of the spatial distribution of the cells. The percolation probability $p$ is 0.45.

**Table. S1** The value and uncertainty of the fitted parameters

| Symbol | Unit | Value±uncertainty |
|---|---|---|
| $\Gamma$ | $s^{-1}cell^{-1}m^2$ | (1.7±0.4)e-5 |
| $\gamma_0$ | $s^{-1}cell^{-1}$ | (2.7±0.09)e-6 |
| $\gamma_1$ | $s^{-1}cell^{-1}$ | (3.3±0.04)e-6 |
| $\epsilon$ | $s^{-1}cell^{-1}$ | (7.4±4.9)e-8 |
| $\epsilon_1$ | $s^{-1}cell^{-1}$ | (4.6±1.4)e-4 |
| $\mu$ | $s^{-1}cell^{-1}$ | (1.8±0.05)e-5 |
| $\mu_1$ | $s^{-1}cell^{-1}$ | (4.9±0.4)e-5 |
| $\lambda_1^{(j,k]}$ (Assuming$\lambda_1^{(j,k]}$ =$\lambda_2^{(j,k]}$ ) | $m^2s^{-1}\ cell^{-1}$ | (1.3±0.4)e-12 |
| $\lambda_2^{(j,k]}$ (Assuming$\lambda_1^{(j,k]}$ =$\lambda_2^{(j,k]}$ ) | $m^2s^{-1}\ cell^{-1}$ | (1.3±0.4)e-12 |
| $\lambda_3^{(j,k]}$ | $m^2s^{-1}\ cell^{-1}$ | (3.5±1.4)e-12 |
| $\lambda_4^{(j,k]}$ | $m^2s^{-1}\ cell^{-1}$ | (4±1)e-10 |
| $\rho_1$ | | 0.75±0.49 |
| $\rho_2$ | | 0.70±0.54 |

### S3 Introduction of Max Permitted Scale (MPS) for sufficient collapse quality

**Inference 1**: If

$$R\left(t|p_j\right) \to R\left(\infty|p_j\right) \equiv \ Const,\ t \to \infty, \tag{S.1}$$

then

$$R\left(T|p\right) \sim p \quad \to \quad R\left(\infty|p\right) \sim p \tag{S.2}$$

where $R\left(\infty|p_j\right)$ is the limit that $R\left(t|p_j\right)$ approaches when $t \to \infty$. Following this, if the $R\left(T|p\right) \sim p$ curves overlap with $R\left(\infty|p\right) \sim p$ as $t \to \infty$, then the scaled curves $RT^{-\zeta/\nu} \sim \widetilde{f}\left(T^{1/\nu}\left(p-p_c\right)\right)$ will be a series of curves moving downright as $T$ increases. The scaled curves are fanned out in such a way that they obviously have not been collapsed. Such scenario is shown in Fig. S2b. Fig. S2b shows the data collapsed by SRCCO, suggesting failure in collapsing the curves for $T = 637.0$h, 713.8h, and 800h.

To collapse the curves with SRCCO (see section 5.7), curves that could lead to failure in SRCCO should be eliminated. Therefore, it is paramount that an upper limit should be put on the choices of the time scales $T$. Here this upper limit is denoted as $T_{max}$

At first the intuition might be observing the time when the derivative of $R\left(t|0.55\right)$ decreases and cut the $T$ value from there, because it is suggested intuitively that the trend in Inference 1 could be avoided to quite a great degree in this way.

Based on the intuition above, a following definition of $T_{max}$ could be proposed: a time scale that is called the Max Permitted Scale $T_{max}$ satisfies

$$\frac{d^2R(t|0.55)}{dt^2}|_{t=T_{max}=0}. \tag{S.3}$$

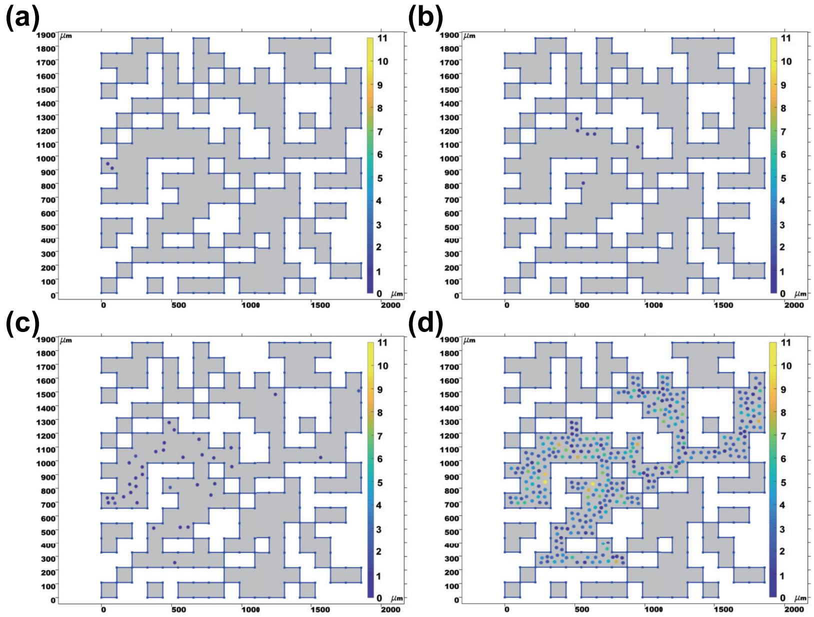


**Fig. S1** An example of the evolution of the spatial distribution of the cells. Cell number here is the number of all three species $S_1$, $S_2$, $S_3$. Figure a-d correspond to the time points $t = 0h$, $t = 204h$, $t = 408h$, $t = 800h$, respectively. Each dot is a node of the triangular FEM mesh. Nodes whose cell number is zero was hidden. The color of the dots represents the cell number corresponding to the node. The label of the color bar is also the cell number.

The optimal critical value set could then be obtained by eliminating the curves with $T > T_{max}$ and then by using SRCCO.

For the model fitted to the raw data from Johnson et al (2019), $T_{max} = 356.3h$ is the solution to equation (S.3). The optimal time scale set $T$ for the curves to be collapsed might be prepared in such a way as

$$\begin{aligned} \mathbb{T}_{optimal}(T_{min},\ \ T_{max}) \equiv \\ &\{T_{min}, \\ &exp\left(log\left(T_{min}\right) + 0.1\sigma\right), \\ &exp\left(log\left(T_{min}\right) + 0.2\sigma\right), \\ &exp\left(log\left(T_{min}\right) 0.3\sigma\right), \\ &..., \\ &exp\left(log\left(T_{min}\right) + 0.9\sigma\right), \\ &T_{max}\}, \end{aligned} \tag{S.4}$$

where,$\sigma = \left(log\left(T_{max}\right) - log\left(T_{min}\right)\right)/10$,$T_{max} = 356.3h, T_{min} = 256h$.

The parameter set derived with $T_{optimal}(T_{min},\ \ T_{max})$ is denoted as $\phi_{optimal}(T_{min},\ \ T_{max})$. Shown in Fig. S2a in the main article are the $R\left(T|p\right) \sim p$ curves with $T \in T_{optimal}(256,\ 356.3)$. Quality of SRCCO could be inferred from the comparison between Fig. S2a and Fig. S2b. It suggests that SRCCO has succeeded in collapsing the curves fanning out in Fig. S2a.

### S4 The FTS result for curves $R\left(T|p\right) \sim p$ with $T \in T_{nolimit}(256)$deviates from that for curves $R\left(T|p\right) \sim p$ with $T \in T_{optimal}(256, 356.3)$

To verify inference 1, the $R\left(T|p\right) \sim p$ curves of the fitted model were scaled into $RT^{-\zeta/\nu} \sim \widetilde{f}\left(T^{1/\nu}\left(p-p_c\right)\right)$ curves using $\Phi_{optimal}\left(256, 356, 3\right)$. Comparison between two types of scaled curves was carried out. The grey curve in Fig. S3b represents the extrapolation of the collapsed curves using $T_{optimal}\left(256, 356, 3\right)$ defined in equation (S.4). And the other curves in Fig. S3b represent the collapsed curves using $T_{nolimit}\left(256\right)$ defined in equation (S.4). It is shown in Fig. S3b in the main article that those curves are fanning out and lower than the grey curve, suggesting failure of FTS when the collapse procedure SRCCO considers the time scales $T > T_{max}$.

### S5 FTS results of the variations

### S6 An illustration of the percolation mesh design

Shown in Fig. S6 is an illustration of the percolation mesh design. A comparison between a sparse mesh and a dense mesh was carried out here. The size of a percolation site should be large enough to avoid the failure illustrated in Fig. S6.

### S7 Slowing down migration does little help to increase the ratio $\frac{T_{max}}{T_{min}}$

Quality of the FTS might be improved by increasing the ratio $\frac{T_{max}}{T_{min}}$. Here is a proposal on how to increase the ratio $\frac{T_{max}}{T_{min}}$ – slowing down the migration to increase $T_{max}$.

Shown in Fig. S7 is the curve of $R\left(t|0.45\right)$, the border-crossing propensity coefficients $\lambda_1$, $\lambda_2$, $\lambda_3$ are all decreased by 256 folds. At time point $t = 0$, $R\left(t|0.45\right) = 0$, and $t_1$ denotes the time point when $R\left(t|0.45\right)$ turns non-zero. $t_2$ denotes the time point when $R\left(t|0.45\right)$ stops changing with $t$. $\Delta t = t_2 - t_1 = 5812h$, $\frac{t2}{t1} \approx 1.9$. The two time points are still not quite separated in logarithmic space. Therefore, Slowing down migration does little help to increase the ratio $\frac{T_{max}}{T_{min}}$.

At the first sight, increasing the size of the percolation lattices is in essence the same as slowing down migration. So, it might not help, either. However, increasing the size of the percolation lattices may serve to enlarge the configuration space, and this might help.

Before moving ahead, let's get familiar with the following definition

$$C\left(t|p_j,k\right) \equiv < B_{jk}\left(t\right) > \tag{S.5}$$

, which is the ensemble mean of $B_{jk}\left(t\right)$.

In fact, as shown in Fig. S8, there were only five conductive ones out of 1000 percolation configurations generated to calculate $C\left(t|0.45,k\right)$,$k=1,2,...,1000$.

From equations (13) and (14) in the main article, it is obtained that

$$R\left(t|p_j\right) \ = \ \left(\sum_{k=1}^{1000} C\left(t|p_j,k\right)\right)/1000 \tag{S.6}$$

Therefore, $t_1$ is the time point when event "$C\left(t|p_j,k\right)$ turning non-zero" occurs for the first time, and $t_2$ is the time point when the event "$C\left(t|p_j,k\right)$ stops changing" occurs for the last time.

So $\frac{t2}{t1}$ may increase if the difference between the configurations is greater.

The typical time of one-dimensional random walk behaves in the following way:

$$\tau_{\mathrm{ty}} = \frac{(\Delta x)^2}{6D} \tag{S.7}$$

where $\tau_{\mathrm{ty}}$ is the typical time, $\Delta x$ the displacement, and $D$ the diffusion coefficient.

The total displacement of the route in Fig. S9a is denoted as $\Delta x_{\mathrm{shortcut}}$, and that in Fig. S9b $\Delta x_{\mathrm{detour}}$. Suppose the size of the percolation lattice in Fig. S9 is $N \times N$.

Obviously,

$$\Delta x_{\mathrm{detour}} \approx \frac{N\Delta x_{\mathrm{shortcut}}}{2} \tag{S.8}$$

And from equation (S.7), the ratio of the typical time of the two routes in Fig. S9, detour to shortcut, is

$$Ratio(\tau_{ty}) = [Ratio(\Delta x)]^2 \ \sim \ o(N^2) \tag{S.9}$$

As $N \to \infty$, $Ratio(\tau_{ty}) \to \infty$, and this suggests a potential surge of $\frac{t2}{t1}$. This possibility still needs exploring in the future.

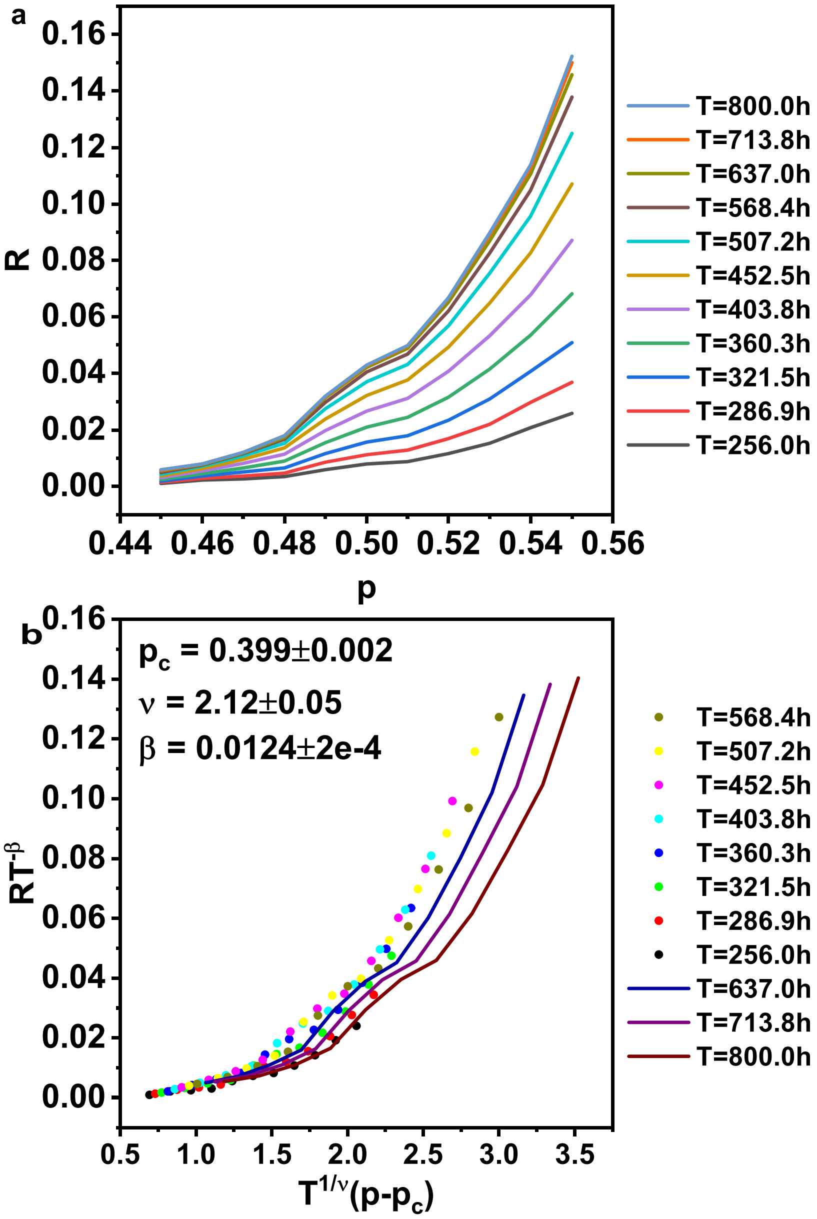


**Fig. S2** Figure a shows the relationship between $p$ and $R(T|p)$. Figure b is the SRCCO result. The critical values $p_c$,$\nu$,$\beta$ were generated by feeding the SPR data $R_{ij}, i \in p, T_i \in T_{nolimit}(256)$. The discrete points correspond to $R_{ij},\ p_j \in p, T_i \in T_{nolimit}(256) \wedge T_i < 637.0$. And the curves correspond to $R_{ij}, i \in p, T_i \in T_{nolimit}(256) \wedge T_i \geq 637.0$.

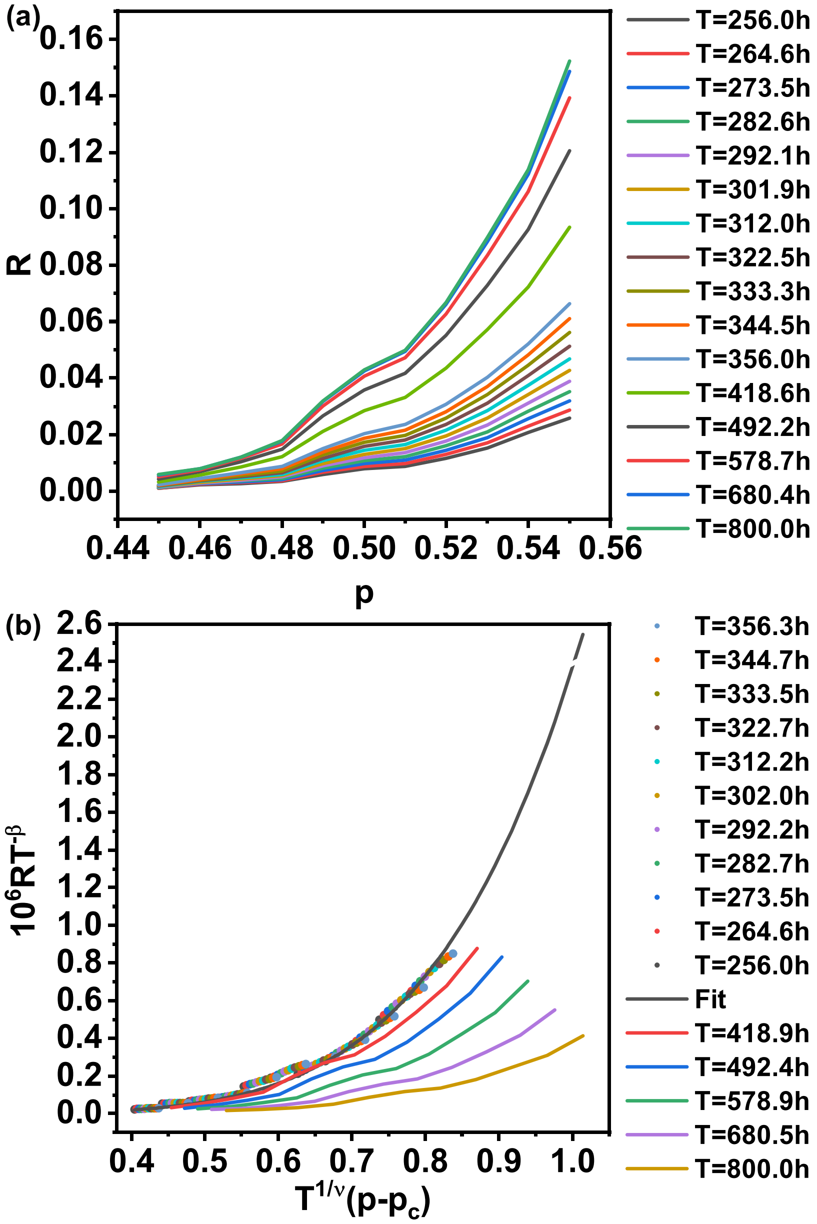


**Fig. S3** Figure a shows the relationship between $p$ and $R(T|p)$, $T_i$ listed on the right. The discrete dots in Figure b are the scaled data $R_{ij}T_i^{-\beta}, p_j \in p, T_i \in T_{optimal}$. The grey curve represents the extrapolation of the discrete dots. The other curves represent the scaled curves as a result of the FTS with parameter $\Phi_{optimal}(256, 356, 3)$, $T_i \in \{418.9,\ 492.4,\ 578.9,\ 680.5,\ 800\}$.

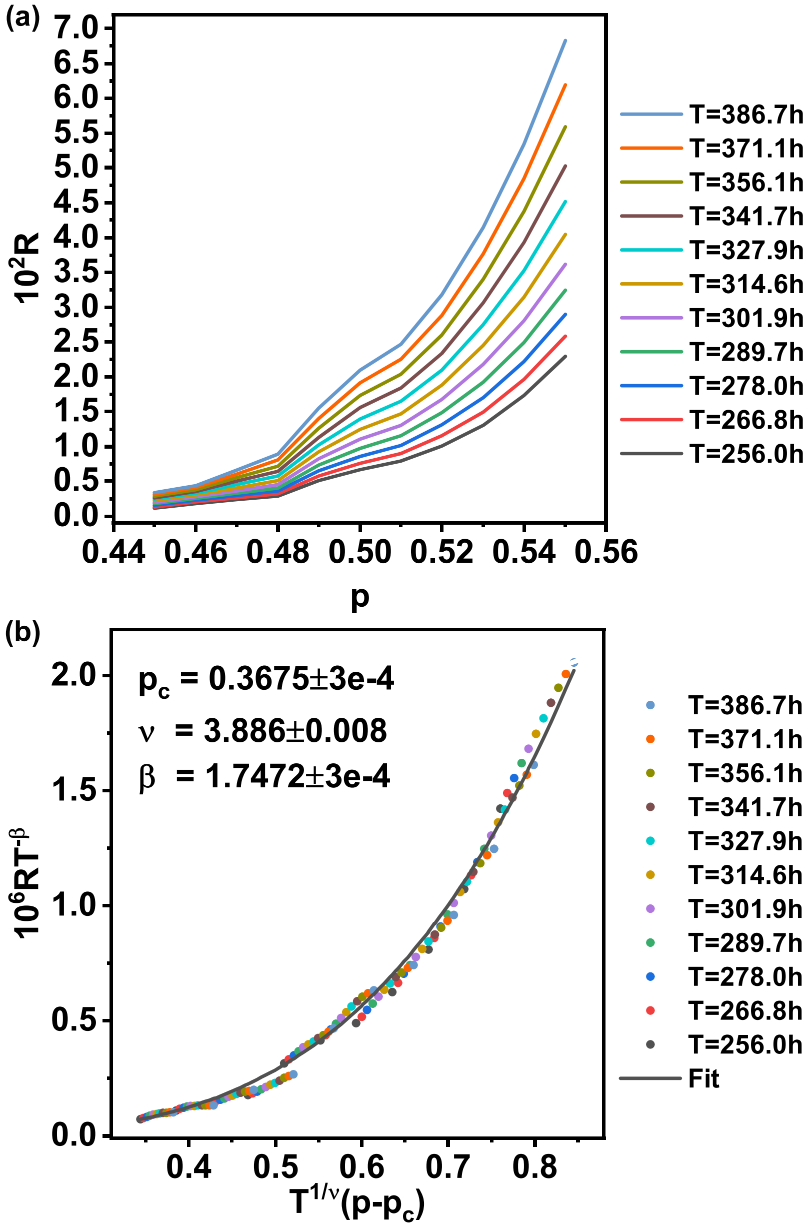


**Fig. S4** The FTS procedure of the tuned model with $\mu_1 = 0$ . Figure a shows the relationship between $p$ and $R\,(T|p)$, $T_i$ listed on the right. Figure b is the FTS result with the critical values $p_c$,$\nu$,$\beta$ generated by SRCCO fed with the SPR data $R_{ij}, p_j \in p, T_i$ listed on the right. The discrete points correspond to $R_{ij}T_i^{-\beta}, p_j \in p, T_i$ listed on the right. The grey curve is the fit for those discrete data.

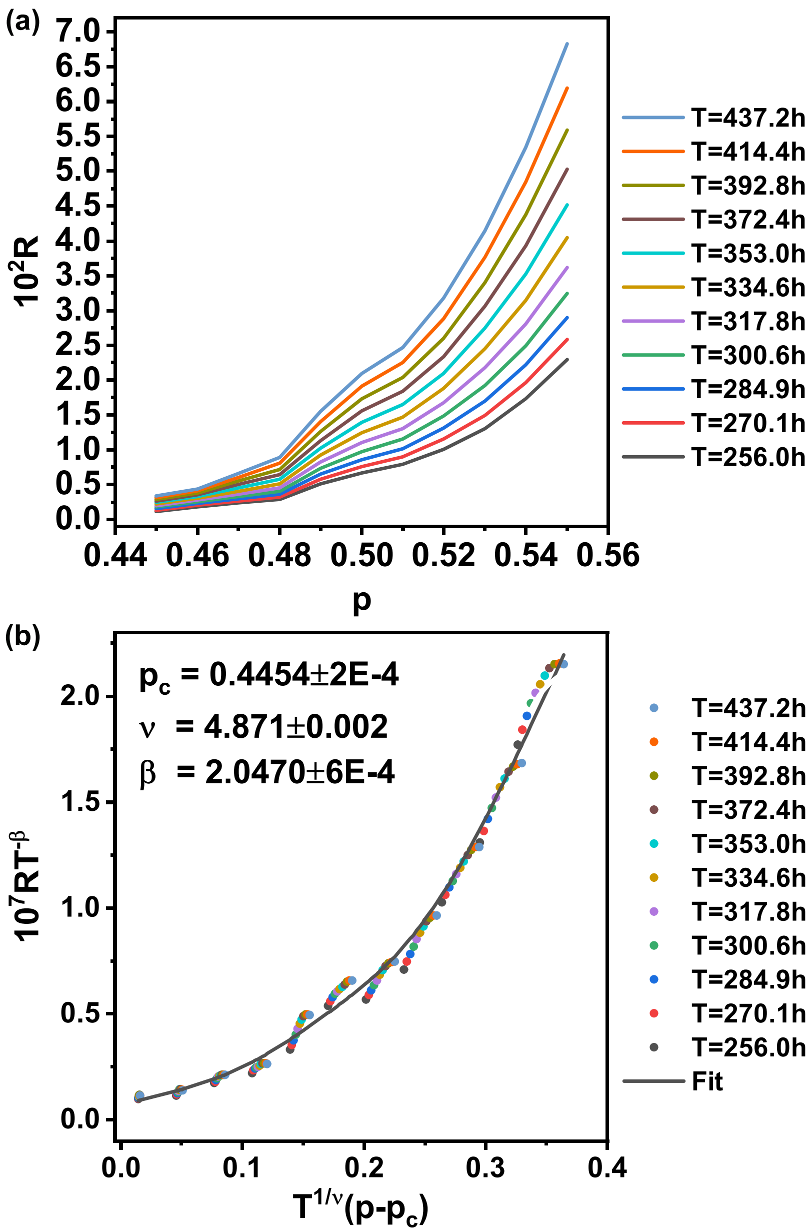


**Fig. S5** The FTS procedure of the tuned model with $\mu_1 = 0$, $\epsilon = 10\epsilon_{original}$ . Figure a shows the relationship between $p$ and $R(T|p)$, $T_i$ listed on the right. Figure b is the FTS result with the critical values $p_c$,$\nu$,$\beta$ generated by SRCCO fed with the SPR data $R_{ij}, p_j \in p, T_i$ listed on the right. The discrete points correspond to $R_{ij}T_i^{-\beta}, p_j \in p, T_i$ listed on the right. The grey curve is the fit for those discrete data.

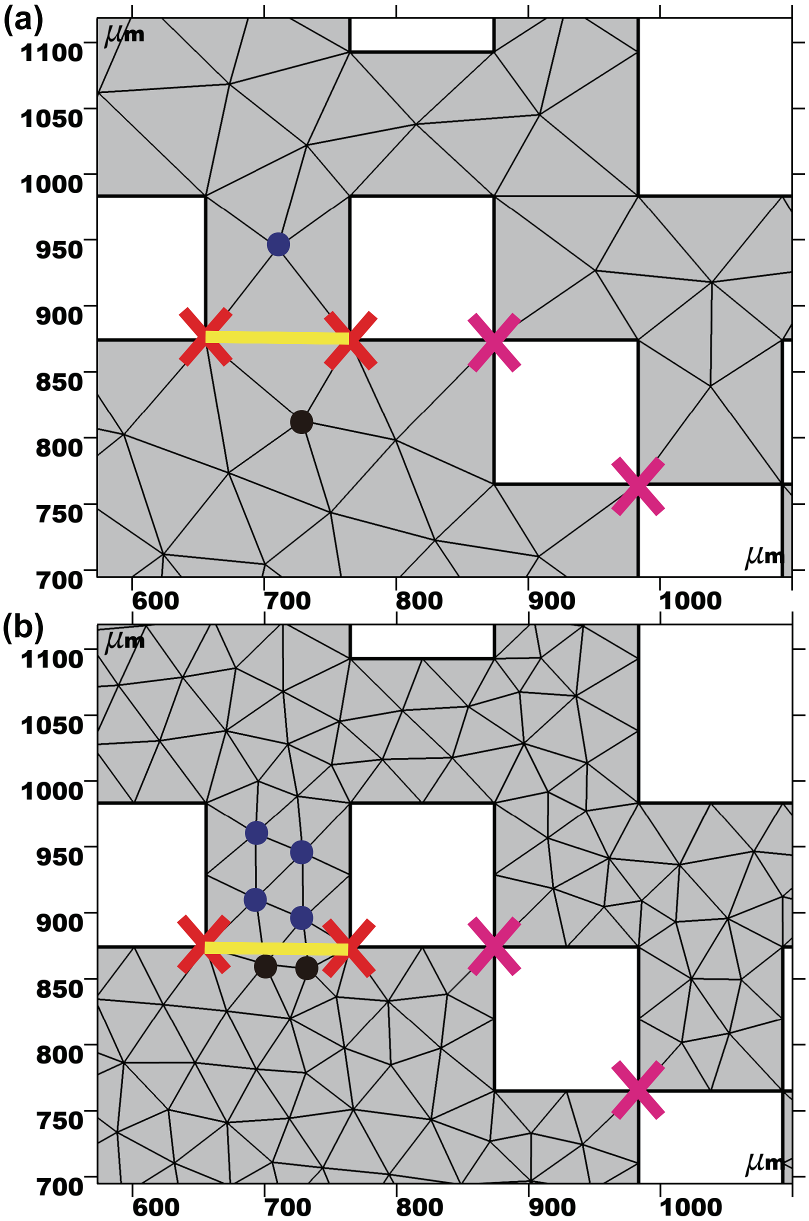


**Fig. S6** A comparison between a sparse mesh and a dense mesh. The crosses are a few examples of the nodes with denied access. The blue dots are all the non-border nodes within the percolation site. The black dots are some nodes in the site below. The magenta crosses emphasize the blocking of border-crossing across the corners. The red crosses represent some of the adjacent nodes of the blue dots. The yellow lines distinct the site above it from the site below. These two sites are connected, though. In figure a, a cell at the blue node can not cross the yellow line because the red crosses are the only options if it is going downwards. Whereas in figure b, a cell at one of the blue nodes can cross the yellow line because the red crosses are not the only options.

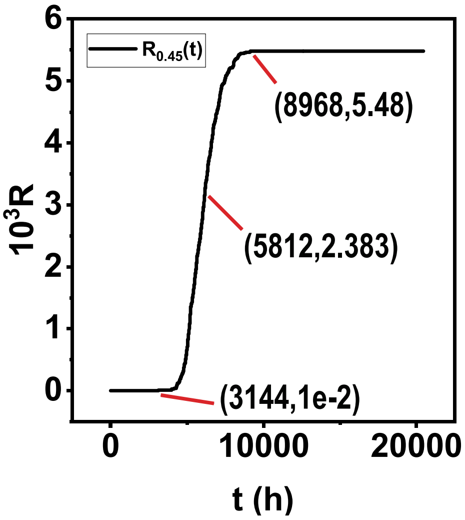


**Fig. S7** The curve of $R(t|0.45) \sim t$, with $\lambda_1$, $\lambda_2$, $\lambda_3$ all decreased by 256 folds. Labels from left to right correspond to the time point when $R(t|0.45)$ turns non-zero, the time point when $R''(t|0.45)$ turns zero, and the time point when $R(t|0.45)$ stops changing with $t$, respectively.

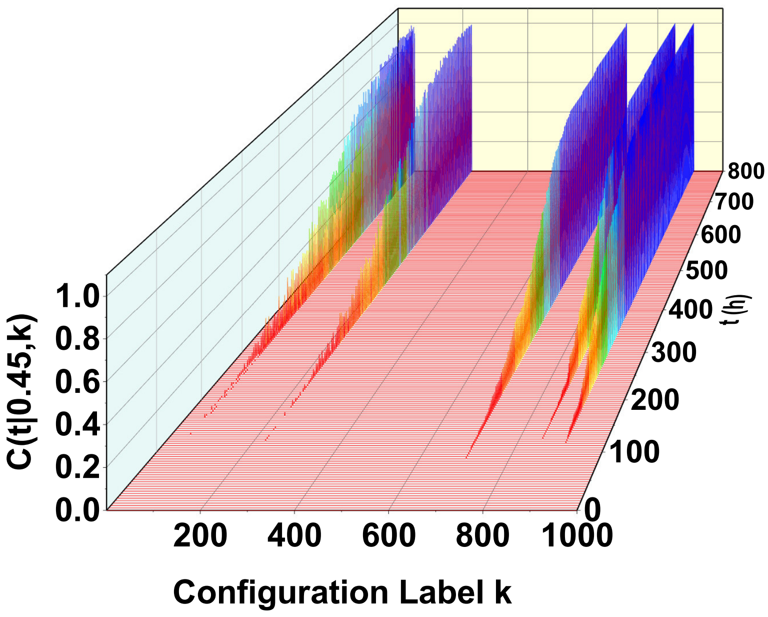


**Fig. S8** $C(t|0.45, k)$, time $t$, and configuration label $k$.

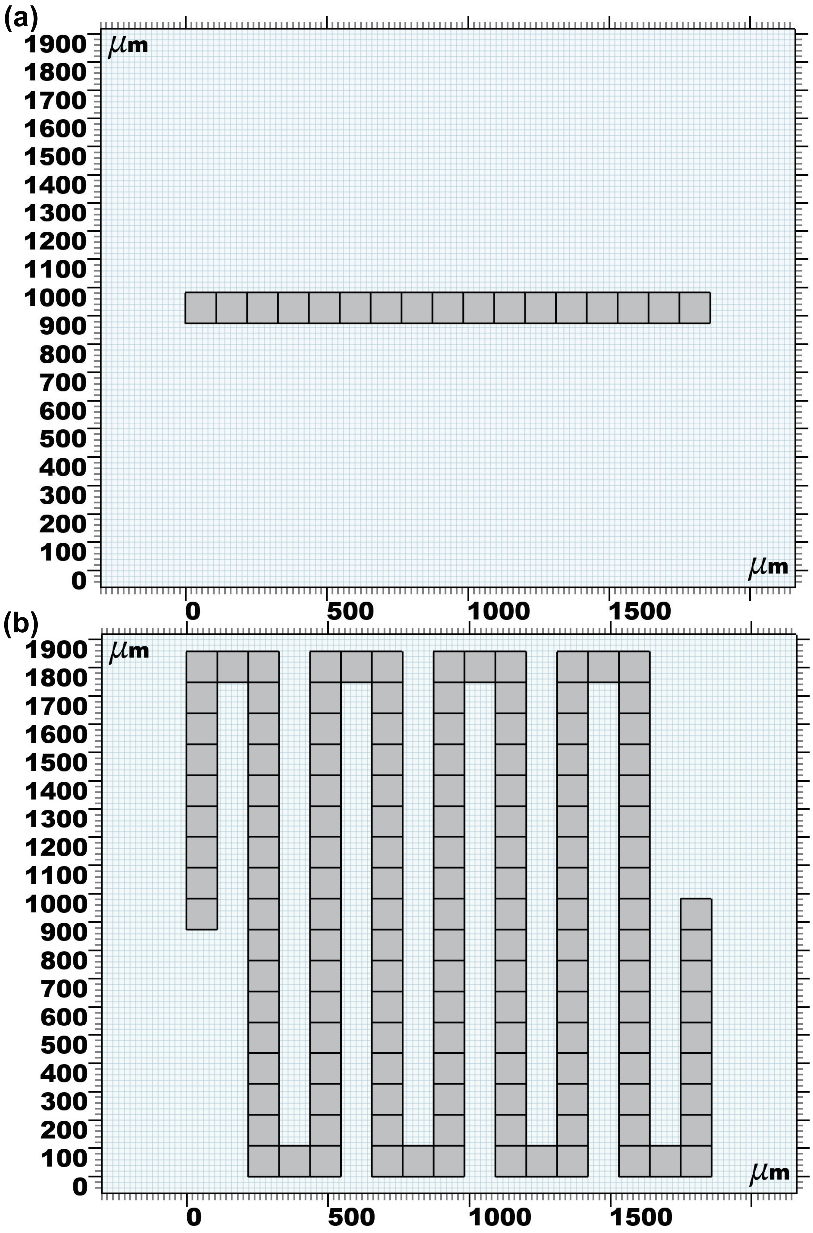


**Fig. S9** Illustration of two extreme types of routes of conductive configurations with the same occupation probability. Figure a represents an extremely short route and Figure b an extremely long one.
